# Monomer-dimer structural comparison in quinol-dependent nitric oxide reductase reveals a functional basis for superior enzymatic activity in the dimer

**DOI:** 10.1101/2024.05.16.593792

**Authors:** Chai C. Gopalasingam, Haruka Egami, Hideki Shigematsu, Masatora Sakaue, Kouki Fukumoto, Christoph Gerle, Masaki Yamamoto, Yoshitsugu Shiro, Kazumasa Muramoto, Takehiko Tosha

**Author notes:** Correspondence (C.C.G), (T.T).

## Abstract

The leading cause of bacterial meningitis, *Neisseria meningitidis,* deploys a quinol-dependent nitric oxide reductase (*Nm*qNOR), belonging to the heme-copper oxidase superfamily. By detoxifying NO, an antimicrobial gas produced by host’s immune system, qNOR enables pathogen survival within hosts. Here, we determined cryoEM structures of the less active monomer and highly active dimer of *Nm*qNOR at resolutions of 2.25 and 1.89 Å, respectively, showing the structural elements responsible for effective NO reduction. Helical disorder at the dimer interface, associated with an altered conformation of the critical Glu563 near the heme/non-heme Fe active site, was observed in the monomer. These findings suggest that dimerization stabilizes the active conformation of Glu563 through the structural network between the dimerization site and the active site. Since other members of the heme-copper oxidases exhibit dimerization, the current data on qNOR helps us understand a regulatory mechanism related to the function of heme-copper oxidases upon oligomerization.

**Teaser:** CryoEM structures unveil a functional rationale for dimerization in nitric oxide detoxifying enzyme from a pathogen

## Introduction

Respiratory enzymes that belong to the heme-copper oxidase (HCO) superfamily play a vital role in cellular respiration under both aerobic and anaerobic conditions. In aerobic respiration, cytochrome *c* oxidase (C*c*O) or quinol oxidase catalyzes a key step for cellular respiration, the oxygen reduction reaction (O_2_ + 4H^+^ + 4e^-^ → 2H_2_O), at a heme/Cu binuclear active center^1^. Conversely, other members of the HCO, cytochrome *c*- dependent nitric oxide reductase (cNOR) and the closely related quinol-dependent enzyme (qNOR), whose active sites consist of heme and non-heme iron (Fe_B_), are crucial for one kind of anaerobic respiration, so-called denitrification^2^. In addition to the critical roles of NORs in anaerobic respiration, catalytic nitric oxide (NO) reduction by NORs (2NO + 2e^-^ + 2H^+^ → N_2_O + H_2_O) is responsible for the elimination of cytotoxic NO produced by hosts’ immune systems in some human pathogens including *Pseudomonas aeruginosa*^3,4^, *Neisseria meningitidis* (*Nm*)^5,6^, and *Staphylococcus aureus*^7^. Given that cellular respiration is an essential biological process for all kingdoms of life and that NORs of pathogens are essential for survival in their hosts, HCO enzymes are a key target for the development of antimicrobial drugs. Indeed, recent work on screening for an allosteric inhibitor for the HCO enzymes demonstrated the great potential for obtaining new antimicrobial reagents^8^. Thus, gathering detailed structural and functional information on the HCO enzymes is imperative for future drug design efforts.

Despite a lower abundance of structural information on NORs compared to the extensively studied C*c*Os, we have conducted several structural studies on NORs and found that qNOR could exist as both a monomer and dimer^9–11^. The biochemical data on qNOR showed that the dimer exhibits 2∼4-fold higher catalytic activity than the monomer in qNOR from *Achromobacter xylosoxidans* (*Ax*) and *Neisseria meningitidis*^10,12^. These observations suggest that the dimer represents the active form in qNOR. Structural analysis with single-particle cryogenic electron microscopy (cryoEM) revealed that transmembrane helix 2 (TM2), an extra helix that is absent in cNOR, assisted the formation of the dimer in qNORs^9–11,13^. However, the distinct reason why the dimer showed higher activity than the monomer in qNOR is not clear, due to a lack of a precise structural comparison of the monomer and the dimer.

Even in the case of other members of the HCO family, like C*c*Os, the relationship of the monomer-dimer states to the function is a principal topic. Recent advances in structural biology provided the fact that C*c*Os usually form supermolecular complexes in the biological membrane^14–20^. For example, human C*c*O is involved in a supermolecular complex consisting of two monomers of complex I (CI_2_), two monomers of complex III (CIII_2_), and two monomers of complex IV (cytochrome *c* oxidase) (CIV_2_)^21^, wholly denoted as CI_2_CIII_2_CIV_2_. However, a supermolecular complex in yeast contains two monomers of complex III and two monomers of C*c*O^18,22^ (CIII_2_CIV_2_) or x1 C*c*O^23^ (CII_2_CIV). It is interesting to note that C*c*O participates in the supermolecular complex exclusively as a monomer, although the crystal structure of the broadly studied bovine C*c*O is a dimer^24^. Furthermore, recent work on the monomer-dimer topic on bovine C*c*O revealed that the oligomeric state is linked to the function of C*c*O, showing that the monomeric form exhibited 2-5-fold higher enzymatic activity than that of the dimer, implying that the monomeric form would be the active form for bovine C*c*O^25^. The structural analysis of each oligomeric state of C*c*Os enriched our understanding of the structure-function relationship in C*c*Os. Therefore, the structural information on the monomer and dimer forms of qNOR will shed light on the molecular mechanism for effective NO reduction in qNOR, leading to further elucidation of the functional importance of constituent oligomeric states of the HCO superfamily.

To that end, we determined the *Nm*qNOR monomer (∼ 85 kDa) structure to 2.25 Å resolution and re-analysed the *Nm*qNOR dimer to 1.89 Å resolution by single-particle cryoEM. The data shed light on how qNOR monomerization deforms a key dimer stabilizing intra-helical arrangement involved in proton transfer and alters the stability of a functionally critical glutamate. These help us further comprehend the structural elements required for an effective catalytic reaction in qNOR, explaining the structural basis for superior enzymatic activity in the dimer.

## Results

### High resolution single particle cryoEM structure of *Nm*qNOR dimer

After purifying and isolating dimer and monomer *Nm*qNOR from the same bacterial culture, we confirmed the enzymatic activity of each oligomeric form and ∼5-fold lower activity in the monomer compared with the dimer (fig. S1). To understand the structural origin for lower enzymatic activity of the monomer than the dimer of *Nm*qNOR, it is essential to carefully compare the structures of the aforementioned oligomeric states. Therefore, we solved the structures of both the monomer and dimer states of *Nm*qNOR using the same cryoEM instrument, a CRYO ARM^TM^ 300^26,27^ (JEOL Ltd.). Owing to the optimization of the conditions for the grid preparation, even though the sample preparation for *Nm*qNOR was fused with apocytochrome *b*_562_ (BRIL) at the C-terminus (details of the sample preparation are in the Methods section), the resolution of *Nm*qNOR dimer was improved to 1.89 Å from ∼3 Å in this study (fig. S2-4, table S1). The high resolution structure of the *Nm*qNOR dimer was essentially the same as our previously determined structure, in which each protomer contains 14 TM helices with heme *b* and the heme *b*_3_/non-heme Fe_B_ binuclear active center, except for the identification of water molecules. The high resolution density map for the *Nm*qNOR dimer allowed us to newly model several water molecules (Fig. 1a). One water cluster is located at the proposed quinol binding site near heme *b*, Asn628, Ser686 and Arg724. Another chain of water molecules is observed around the propionates of heme *b*_3_ and Ca ion as observed in the other qNORs (*Ax* and *Geobacillus stearothermophilus* (*Gs*)) and even in cNOR from *P. aeruginosa*^13^ (fig. S5-6). Notably, only a few water molecules are observed in the presumed water channel from the cytoplasmic side to the active center in the high resolution *Nm*qNOR dimer (Fig. 1b), unlike the similarly high resolution (2.25 Å) cryoEM structure of the *Ax*qNOR dimer^28^ and X-ray crystal structure of *Gs*qNOR^11^. This observation suggests that the apparent lack of discrete water molecules in the putative water channel of the *Nm*qNOR dimer could be due to a more mobile and dynamic nature of the water molecules in the channel for effective catalytic proton transfer.

**Fig. 1.**
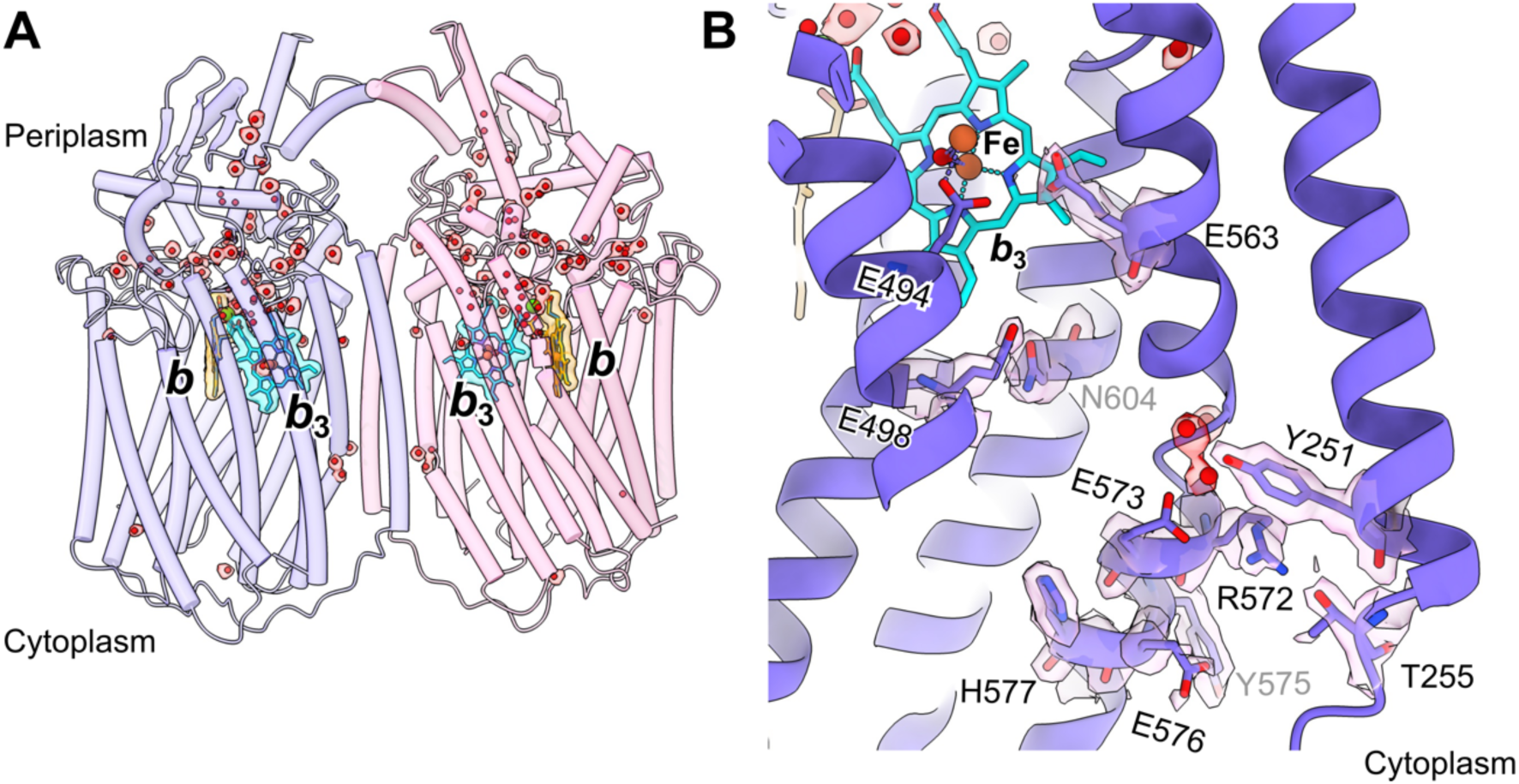
High resolution cryoEM structure of the *Nm*qNOR dimer. (**A**) Cylinder representation of the *Nm*qNOR dimer, with individual protomers colored in slate blue and hot pink. Water molecules are shown as red spheres surrounded by transparent red density map, with heme *b* shown as cyan sticks and heme *b*_3_ as orange sticks. (**B**) View of the potential proton transfer channel from the cytoplasmic end to the active site heme *b*_3_ and non heme iron (brown sphere), with residues shown as blue sticks with surrounding density in transparent pink.

### Single particle cryoEM structure of the *Nm*qNOR monomer

The *Nm*qNOR monomer is ∼ 85 kDa, asymmetric, and lacks large, extracellular domains, making it a relatively challenging target for sub-3 Å cryoEM reconstruction. Nevertheless, we obtained a structure with a resolution of 2.25 Å from 274,346 particles using data collected on a CRYO ARM^TM^ 300 (JEOL Ltd.) and cryoSPARC v4.1.1 for image processing (fig. S7-9, table S1). The density for the BRIL portion is slightly visible, yet no model could be built for it as in the case of the *Nm*qNOR dimer. The overall structure of monomer *Nm*qNOR is almost identical to a protomer of the *Nm*qNOR dimer (Root mean square deviation = 1.35 Å for Cα atoms). Akin to the high resolution dimer structure, the EM map of the monomer revealed the location of water molecules near possible quinol binding site and the propionates of heme *b*_3_, yet, not in the putative water channel from the cytoplasmic side (Fig. 2a,b).

**Fig. 2.**
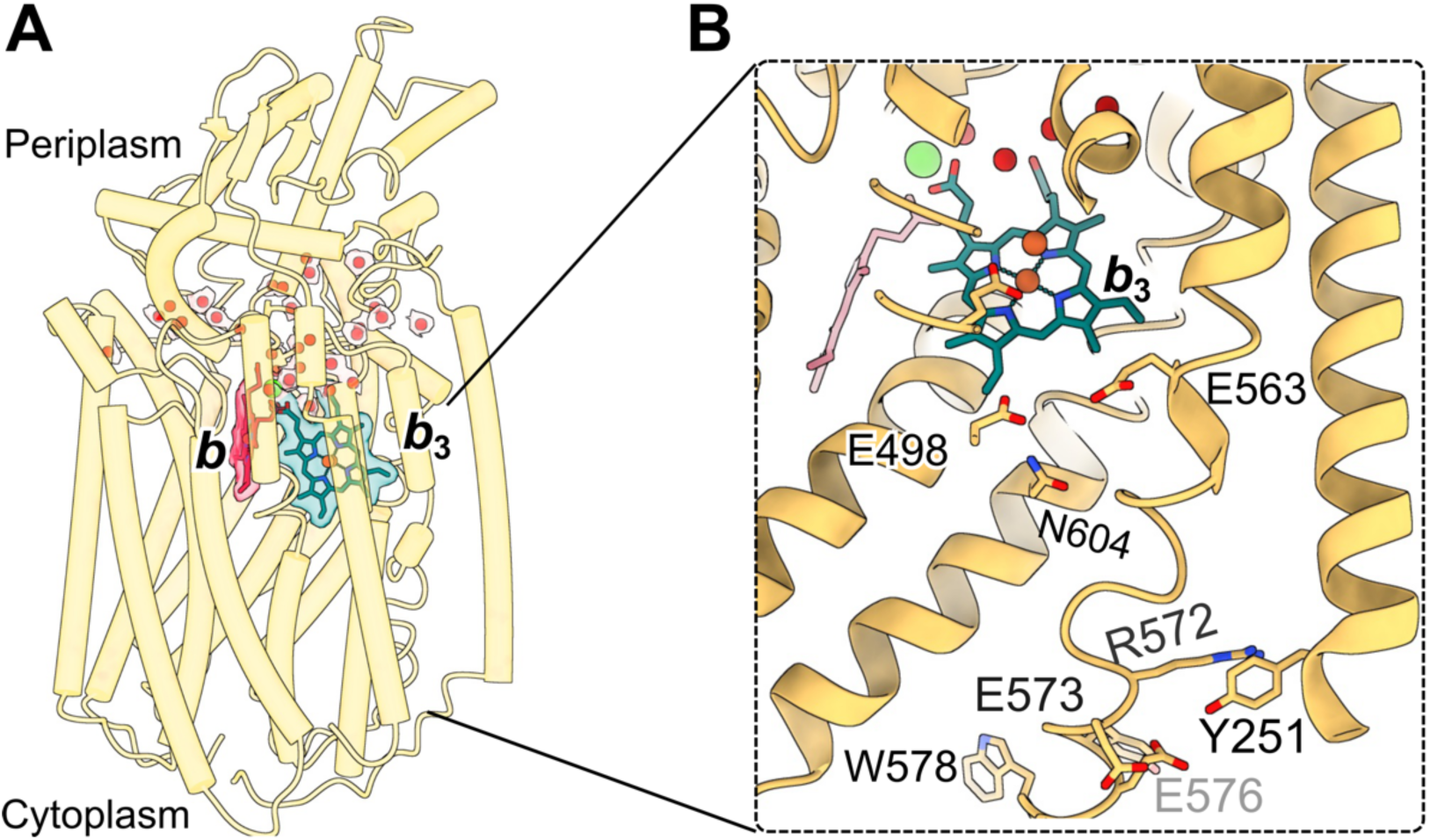
High resolution cryoEM structure of the *Nm*qNOR monomer. (**A**) Cylinder representation of the *Nm*qNOR monomer (gold) with water molecules shown as red spheres surrounded by transparent density, heme *b* as red sticks and heme *b*_3_ as teal sticks. (**B**) View of the potential proton transfer channel in ribbon representation, from the cytoplasmic end to the active site, with residues shown as gold sticks.

### Structural comparison of the *Nm*qNOR monomer and dimer

The reconstructions at comparable resolutions for the monomer (2.25 Å) and the dimer (1.89 Å) of *Nm*qNOR allow us to directly compare the structure of the monomer with that of dimer to get insights into the enzymatic activity difference between the two forms. In particular, the significance of higher resolution reconstructions can divulge changes in water networks, side chains and/or heme propionate conformation. These components are all pivotal to enzyme functionality, with even subtle alterations potentially causing large differences in enzyme reactivity. We found substantial differences both at the active site and the TM helices related to dimerization (and proton transfer) in the monomer and dimer (*vide infra*), although the other structural elements such as, the configuration of the redox centers, putative quinol binding site and the potential hydrophobic NO binding/transport channel remained essentially identical (fig. S10). The putative quinol binding site, which consists of the conserved His303, Glu307 and Asp728, are unchanged in their conformations (fig.S10b-c).

#### I) Heme/non-heme Fe binuclear active center

Figure 3 represents the catalytic site of qNOR, consisting of a high spin heme *b*_3_ and a non-heme iron, Fe_B_, in both the monomer and dimer states. The coordination structure and the geometry of the heme *b*_3_/Fe_B_ active center are comparable in the monomer and the dimer. For example, in addition to three conserved histidine residues (His490, His541, His542), the conserved Glu494 coordinates Fe_B_ in a monodentate manner both in the monomer and the dimer. The distances between heme *b*_3_ iron and Fe_B_ in the monomer and the dimer are 3.64 and 3.9 Å, respectively, both of which are reasonable distances to accommodate the presence of an μ-oxo bridging ligand^29–31^. However, the dynamic property of conserved Glu494 could likely be altered between the monomer and dimer. In the dimer, the density assignable to Glu494 was only clear at the low sigma level (Fig. 3a), indicating the flexible nature of this residue. However, this property of Glu494 was not observed in the monomer active site (Fig. 3b), despite similar reconstruction resolutions and total electron dose applied during data collection.

**Fig. 3.**
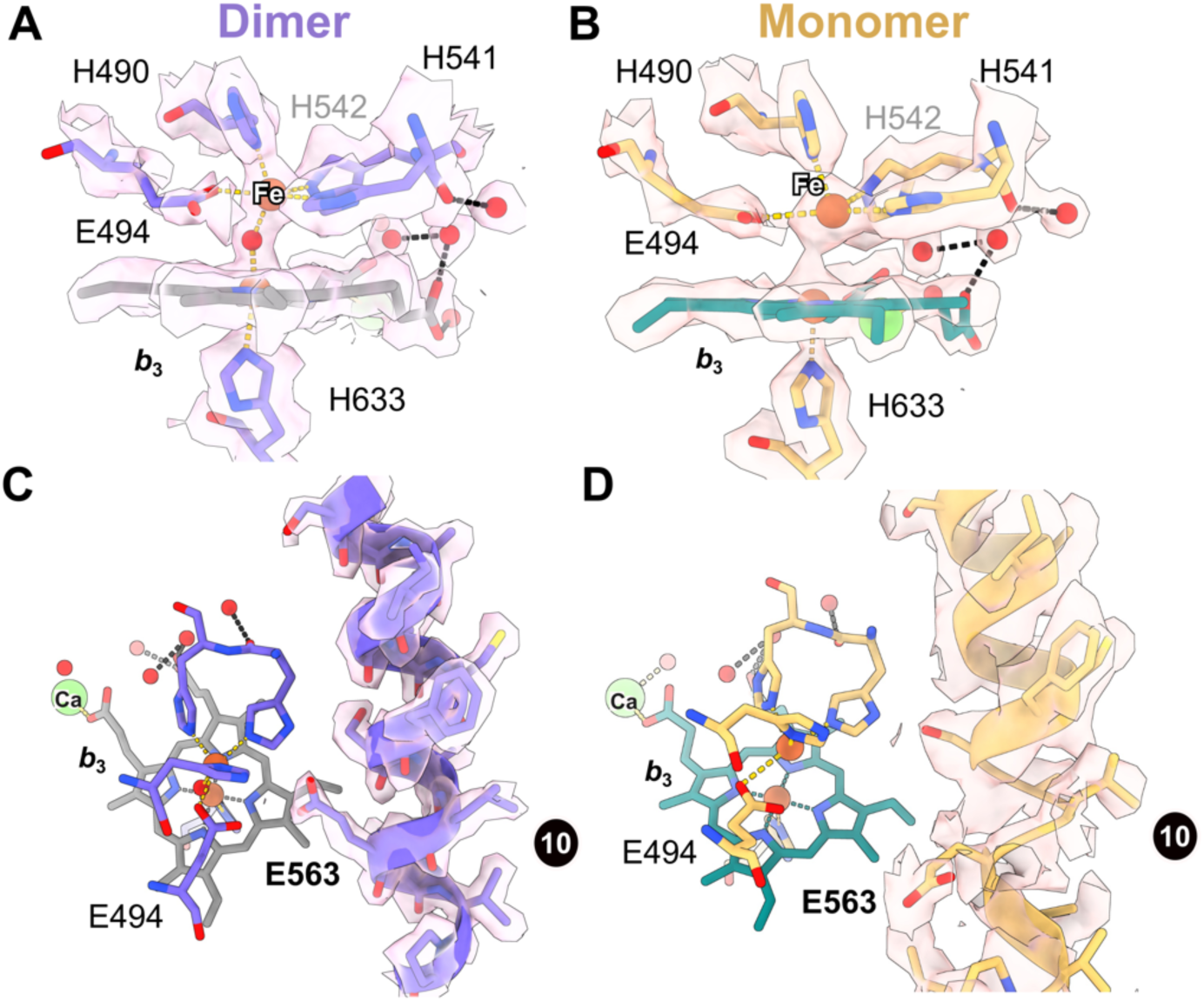
The active site structures of the *Nm*qNOR monomer and dimer. (**A**-**B**) Active site of the dimer (A) and monomer (B) with model and map representation. Metal-ligand bonding is shown with gold dashes, whilst water-ligand bonding is shown with black dashes. (**C-D**) View of TM10 and associated Glu563, and their interaction in the context of the active site for both dimer (C) and monomer (D) structures. Heme *b*_3_ is shown as grey and teal sticks, for the dimer and monomer structures. Non-heme iron and calcium ions are depicted as brown and green spheres, respectively.

In addition to possible changes in the dynamics of Glu494, the orientation of conserved Glu563, in the second coordination sphere for Fe_B_, is different between the monomer and the dimer of *Nm*qNOR (Fig. 3c,d). The carboxylate group of Glu563 is facing towards Fe_B_ and is within hydrogen-bonding distance (3.3 Å) of the carboxylate group of Glu494 in the dimer, whereas the side-chain of Glu563 is flipped away (or, not in a metastable conformation) from Fe_B_ in the monomer. Since Glu563 is suggested to work as a proton donor for catalytic NO reduction at the end of the water channel^32,33^, it is highly plausible that the dynamic, conformational difference of Glu563 is associated with the lowered enzymatic activity of the monomer than that of the dimer.

#### II) TM helices involved in the dimerization

Given that Glu563 is located on TM10, a helix responsible for the dimerization of *Nm*qNOR, it is of great interest to compare the structural properties of TM10 in the monomer and the dimer. The current high resolution structure of the *Nm*qNOR dimer indicated that TM10 is responsible for dimerization (Fig. 4a). Ala574 and Trp578 of TM10 and Val598, Phe602, and Ile606 of TM11 (chain A) effectively sandwich the lower portion of TM2 via interactions with Leu246, Trp249, and Phe253 from TM2 of the other monomer (chain B) enabling dimerization (Fig. 4a). Additionally, Leu569 in TM10 and Ile244 in TM2 of each monomer forms a hydrophobic core to make a four-helix bundle (TM2A, B and TM10A, B) like structure (Fig. 4b). Contrary to this, in the monomer form, TM10 is notably free from any other interactions (Fig. 4c). The lower portion contains residues Tyr575 and Trp578, which help form the dimer interface, and the accompanying density is much broader and less defined than the surrounding density (Fig. 4c). In the absence of the four-helix bundle-like structure, the monomer Leu569 (on TM10) is rotated away from this pocket (Fig. 4d).

**Fig. 4.**
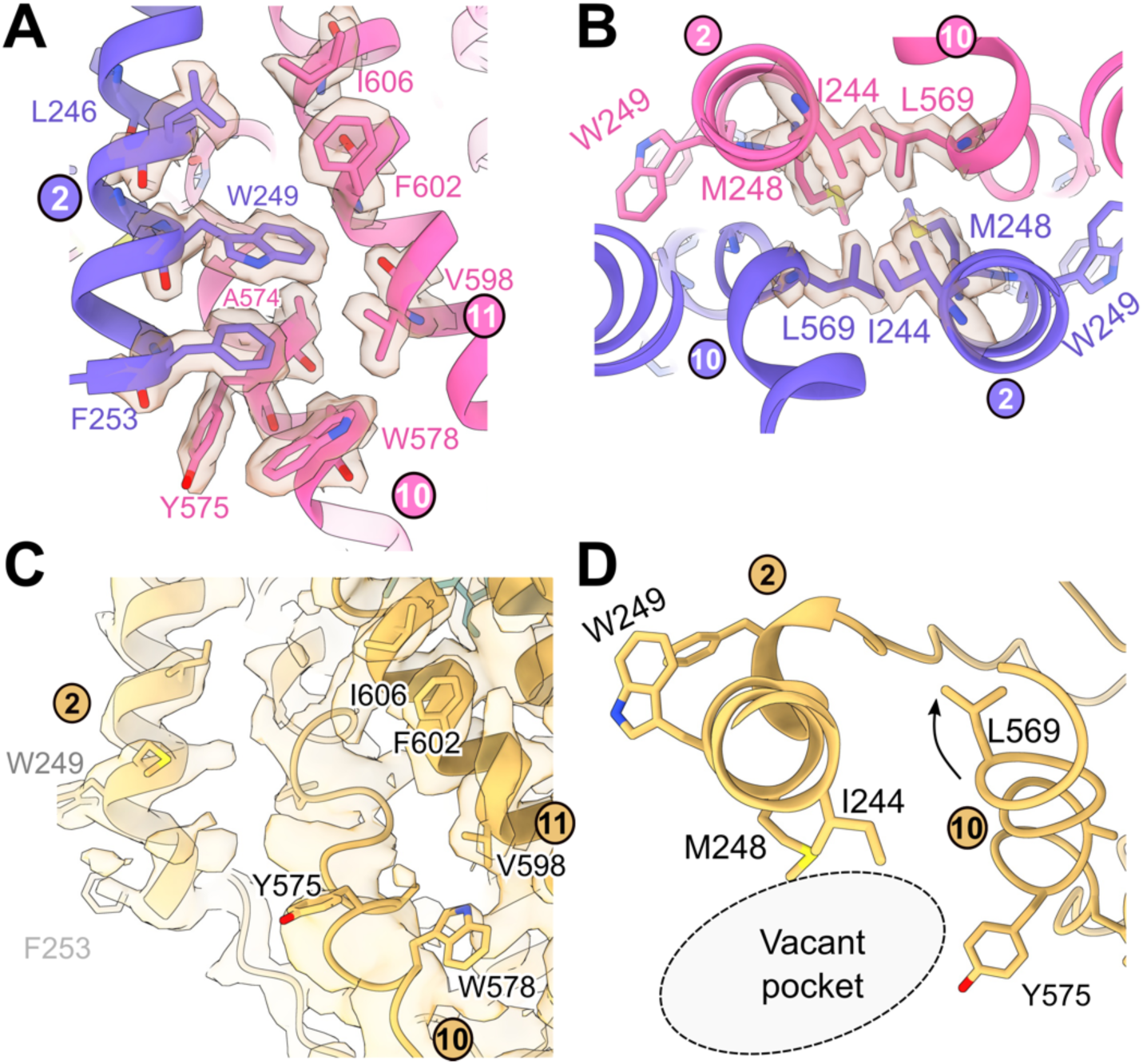
The dimerization sites of *Nm*qNOR monomer and dimer. (**A**) TM2 of one protomer (slate blue) forming intrahelical interactions with TM10 and TM11 of the other protomer (hot pink), mediated by W249, F253 on TM2, F602 on TM11, and W578, and Y575 on TM10. (**B**) View from the periplasmic side showing the qNOR dimerization site; the four-helix bundle-like structure and the hydrophobic residues forming the interaction site. Residues are contoured with the sharpened map densities. (**C**) The monomer structure’s dimerization site shows an absence of TM2 from the other protomer. TM10 displays broader, smeared density compared to the dimer. (**D**) View from the periplasmic side of (C), showing TM10 Leu569 rotation and the vacant pocket of the dimerization site in the monomer.

The above observations suggested that the monomer *Nm*qNOR TM10 may have increased flexibility. To improve the clarity in these regions, we performed focused 3D classification with no alignment and masked refinements. However, this did not yield reconstructions with discrete conformations of the full region of TM10 or TM2. In an attempt to describe the potential, continuous motion of TM10 and 2 within the monomer particle images, we performed 3D Variability Analysis (3DVA) in cryoSPARC v4.4.1^34^, which has been successfully employed in studies of G-protein coupled receptor dynamics^35,36^. As proteins often exhibit some degree of flexibility for their mechanism, there are potentially many possible 3D structures that it can adopt. Instead of reconstructing one single 3D structure from particle images, i.e., assuming one rigid conformation, this algorithm reconstructs a family of related 3D structures. These can help describe the various flexible conformations (subunit rotation/ hinge-like movement, helical bending and swaying etc.,) that are present in the molecule. This provides a visual insight into the conformational space of a macromolecule using 3D movies to record the movements (frames).

After analyzing the *Nm*qNOR monomer with a solvent mask excluding the detergent micelle (to focus only on the protein dynamics), significant swaying and bending of TM10 were clearly observed, with additional minor, lateral movements of TM2 (Movie S1-3). Aside from this, no other portion of the molecule displayed such dynamic motions. Across three different trajectories (variability components) in the space of 3D structures where the molecule exhibits variability, all of them resolved the swaying of TM10 back and forth into the dimer interface pocket, with a 7 Å maximal positional difference between the first and last frame of the 3D movie series (Fig. 5a). The region from Pro566 to Gly571 begins to display greater variability of helical angle and direction (Fig. 5a). Variability component 2 resolved intermediate movements compared to others, with more straightening of the bottom portion of TM10 (Fig. 5a). The conformational flexibility of TM10 in the dimer form of *Nm*qNOR was also analyzed using 3DVA, yet, no such characteristic motion was observed in TM10 of either protomer (Movie S4-6, Fig. 5b), with only minimal, rigid body motion (slight rotation) of the protomers observed. These results indicated that only the monomer TM10 conformational landscape is less restricted (e.g., significant conformational variability), due to the lack of the neighboring protomers TM2 and TM10 interaction to stabilize its conformation.

**Fig. 5.**
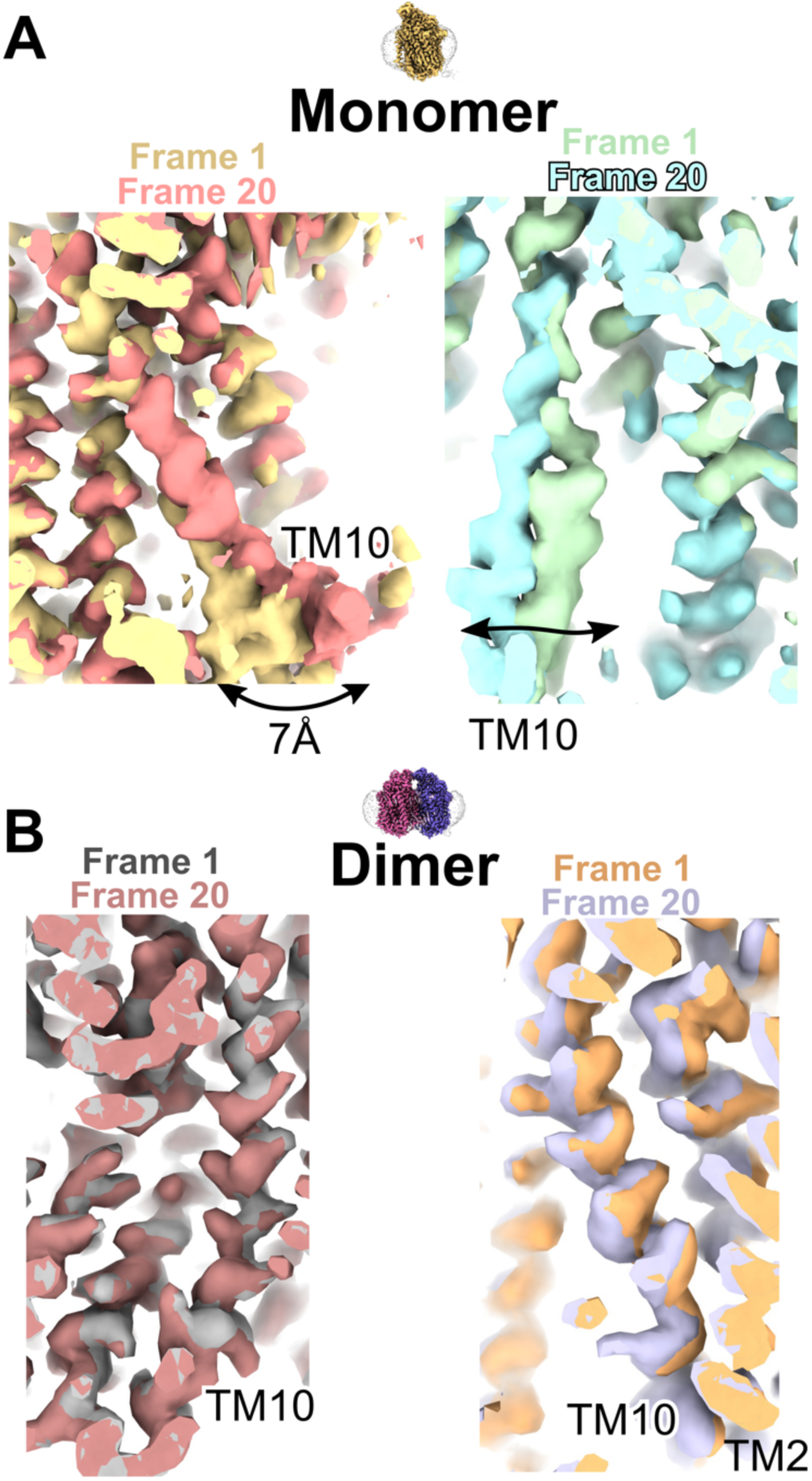
The dynamic property of TM10 in the *Nm*qNOR monomer. 3DVA results from the *Nm*qNOR monomer and dimer. (**A**) Representative snapshots from the 3DVA of the *Nm*qNOR monomer. Superposed frames 1 and 20 from component 1 and 2 are displayed in the left and right columns, respectively. Large differences in TM10 conformation are evident in both component analyses. (**B**) Representative snapshots from the 3DVA of the *Nm*qNOR dimer. Superposed frames 1 and 20 from component 1 and 2 are displayed in the left and right columns, respectively, in the same viewing orientation as shown in panel (A). Frames are almost indistinguishable from each other (comparatively little motion) compared to the monomer TM10 movements.

The difference in the dynamics of TM10 in the monomer and the dimer is illustrated by the modeled conformational and structural change in TM10 compared to the dimer (Fig. 6a). The TM10 helix in the dimer has a kink formed by highly conserved Pro566 and Gly571 residues (Fig. 6a, fig. S11), which creates a 35° bend in the helix. The dissociation of the dimer causes the outward swinging motion (∼ 6 Å) that occurs from Gly571 onwards in TM10. The lower portion of monomer TM10 partially occupies the space where the opposing protomer TM2 exists, in the dimer form and induces a positional shift of TM2 by ∼3.0 Å (Fig. 6a). Although the area around Glu563 in TM10 shows a relatively small positional shift in the monomer as compared with the dimer (Fig. 6b), the broad structural differences of TM10 involved in the dimerization of *Nm*qNOR would be accompanied by an orientation change and increased flexibility of conserved Glu563 between the monomer and the dimer (Fig. 6b).

**Fig. 6.**
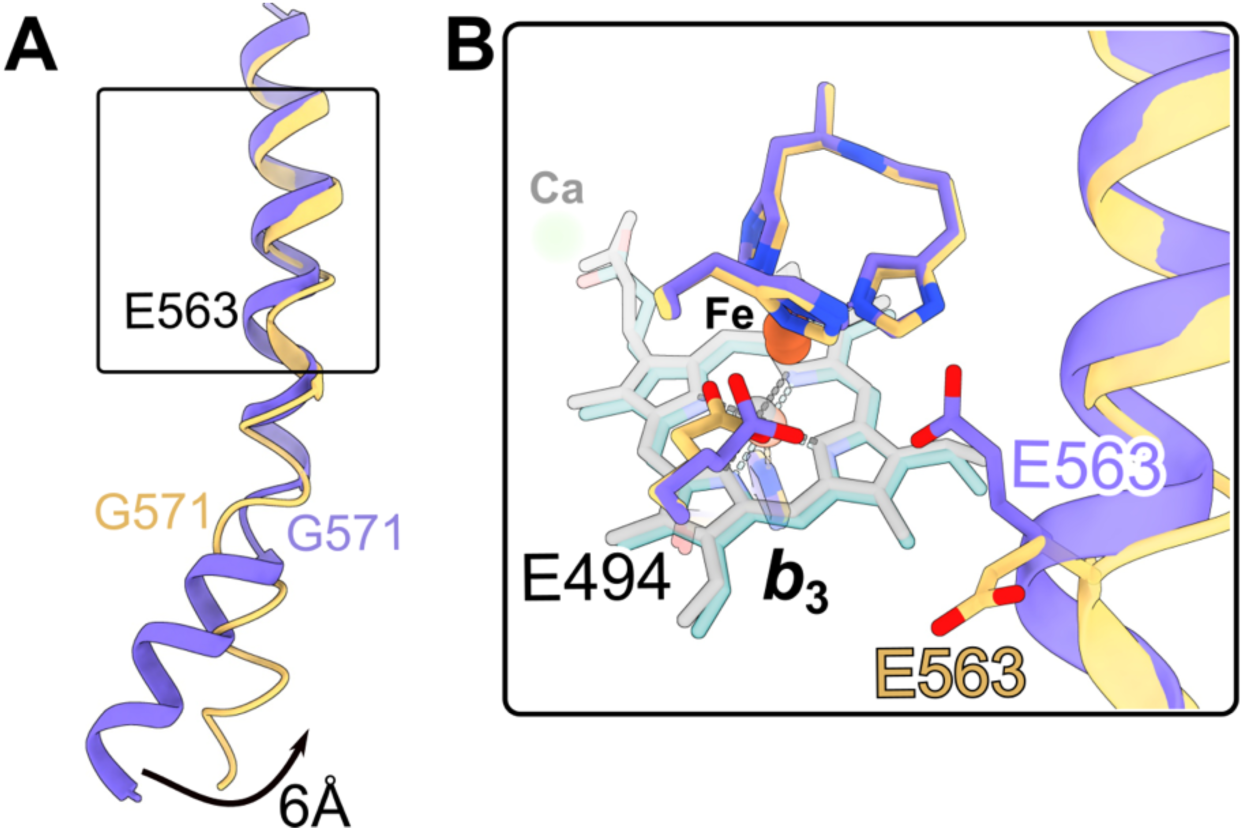
Structural comparison of TM10 in the monomer and dimer *Nm*qNOR. (**A**) Comparison of TM10 between monomer (gold) and dimer forms (blue). Ribbon model representation showing the helical swing (black curved arrow) in the cytoplasmic region, after G571. The black box is shown in detail in panel (B). (**B**) Superposition of the active sites of *Nm*qNOR monomer (gold) and *Nm*qNOR dimer (slate blue), showing differing conformations of Glu563, which is located on TM10. Heme *b*_3_ is shown as transparent gray and teal sticks for the dimer and monomer structures, respectively.

### Structural and functional effects of the mutations of conserved Glu563 in *Nm*qNOR

The substitutions of conserved Glu563 with Ala and Leu, both of which cannot form a hydrogen bond with Glu494, confirm the functional importance of Glu563 and postulate a possible role in maintaining the dimer assembly. As summarized in Table 1, the substitutions of Glu563 lowered the NO reduction activity to less than 10% of the wild-type dimer, even in their dimeric states (Fig.7a). In addition to the crucial role of Glu563 in the enzymatic function, we found that the mutations also affected the dimerization as evident from the profile of the size exclusion chromatography (Fig. 7b). Analysis of the oligomeric content by blue native PAGE showed that both mutations of Glu563 increased the population of the monomer as compared with wild-type (Fig. 7c). Thus, Glu563 is suggested to be essential for the enzymatic function, and to be important for the dimer formation through the interaction with Glu494.

**Fig. 7.**
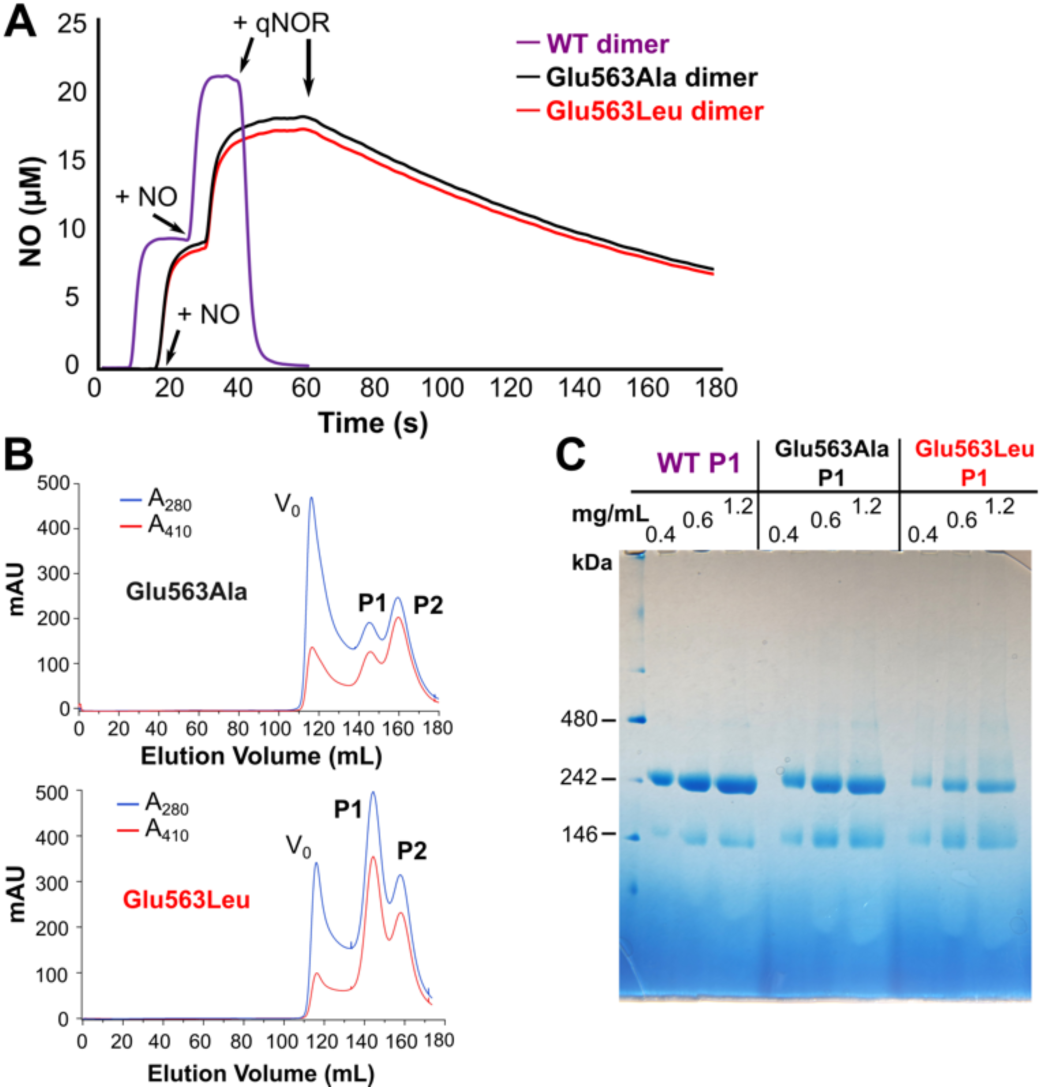
Functional and native PAGE analysis of wildtype and Glu563 variants of *Nm*qNOR. **(A)** NO reduction activity measurements (representative traces) of wildtype (WT) dimer (purple trace), Glu563Ala (black trace), and Glu563Leu (red trace) dimer samples. The addition of substrate NO and qNOR enzyme are indicated by arrows. (**B**) Size exclusion chromatograms of Glu563Ala (top) and Glu563Leu (bottom). V_0_, P1, and P2 refer to column void volume, peak 1 and peak 2 elution positions. Blue and red traces correspond to absorbance values at 280 and 410 nm, respectively. (**C**) Blue native-PAGE results of wildtype peak 1 (dimer) samples against Glu563Ala and Glu563Leu peak 1 sample. 0.4, 0.6 and 1.2 indicated the final, loaded concentrations in mg/mL for each lane. The molecular weight ladder is shown on the left with labels.

**Table 1.** The effects of the substitutions of Glu563 on the catalytic activity of *Nm*qNOR.

| Protein | NO consumption rate<br>($\mu\text{mol-NO s}^{-1}$ per $\mu\text{mol}$<br>qNOR) | Sampling |
| --- | --- | --- |
| **Wildtype** |  |  |
| Dimer | $79 \pm 6$ | 7 biological replicates, 5<br>technical replicates |
| Monomer | $17 \pm 5$ | |
| Monomer (for cryoEM<br>analysis) | $16 \pm 7$ | 4 biological replicates, 5<br>technical replicates |
| **Mutants** |  |  |
| Glu563Ala Peak 1 | $1.6 \pm 0.4$ | 5 technical replicates |
| Glu563Leu Peak 1 | $0.5 \pm 0.2$ | |

## Discussion

In this study, we obtained high-resolution cryoEM structures of the monomer (2.25 Å resolution) and the dimer (1.89 Å resolution) states of *Nm*qNOR from the same batch of bacterial culture. The attained resolution of the dimer is the highest resolution for any NOR structure thus far, and generally one of the highest resolution single particle reconstructions for a macromolecule of its molecular weight range (< 200 kDa). Having similar resolution range structures determined with the same cryoEM instrument allows us to perform a detailed structural comparison of the monomer with that of the dimer, to get structural insights into the origin of superior enzymatic activity in the dimer over the monomer.

As the NO reduction reaction by qNOR requires three elemental processes; substrate NO binding, electron transfer from quinol, and proton transfer, the lowered enzymatic activity in the monomer form can be caused by suppression of at least one of the aforementioned processes. As mentioned in the results section, the possible NO binding channel, a Y-shaped hydrophobic channel in the TM region^37^, is almost indistinguishable between the monomer and the dimer of *Nm*qNOR (fig. S10d-f), suggesting that the substrate binding process could be independent of the oligomeric state of the enzyme. In addition, the presumed quinol binding site, which is composed of conserved His303, and Asp728, is not majorly affected by the dimer formation in *Nm*qNOR (fig. S10b,c). This suggests that the property of quinol binding is not a factor affected by dimerization. Furthermore, the redox potentials of hemes, which are related to the rate of electron transfer, are unlikely altered by the dimerization, since the water network around the active site and heme propionates, determinants of the redox potential^38–40^, are almost identical between the monomer and the dimer (fig. S10a). It is, therefore, less possible that the electron transfer process is altered by the dimerization of *Nm*qNOR.

It was proposed from several structural and structure-based mutational studies that protons required for the catalytic reaction are supplied from the cytoplasm through a putative water channel in qNOR^11,12,41^. There is a hydrophilic large cavity connecting the cytoplasm and the active site in the current cryoEM structures of both monomer and dimer, indicating that a possible water channel could be conserved in both states. However, the region around the active site portion of the hydrophilic channel (i.e., the terminal point of said channel) exhibits marked difference between the monomer and the dimer. The side-chain of Glu563, one of the conserved Glu residues near the binuclear active center, is towards the Fe_B_ ligand, Glu494, in the dimer, whereas the side-chain and is flipped away from the active site in the monomer (Fig.3). Amongst a selection of NOR structures, only the X-ray crystal structure of *Nm*qNOR with zinc bound displays a similar Glu563 conformation^10^ as we observed in the current cryoEM monomer *Nm*qNOR structure (fig. S12). The Zn-bound structure likely represented an inactive state, based on in-solution activity measurements^10^. From the current study, we further strengthen the notion that Glu563 is essential for the enzymatic activity of qNOR (table 1). The orientation and dynamic variation of the side chain would alter the efficiency of essential proton transfer, thereby leading to lowered NO reduction activity in the monomer.

Given that Glu494 is a possible terminal proton donor for the catalytic NO reduction^41–43^, increased structural dynamics of Glu494 upon dimerization would also contribute to the enhancement of the catalytic activity. The presence of the carboxylate group of Glu563 close to the carboxylate group of Glu494 might increase the flexibility of Glu494 via electrostatic repulsion, thereby heightening catalytic activity in the dimer. Thus, dimerization can manipulate the structure around the end of the active site side of the water channel for effective proton transfer in the catalytic NO reduction reaction.

The here presented structural data provides a rationale concerning the mechanism of the observed orientation change of conserved Glu563 upon the dimerization of *Nm*qNOR. Comparison of the respective oligomeric states reveals that dimerization induces kink formation of TM2 and TM10 through intra-protomer interactions of TM10 and TM2, respectively (Fig. 4). Furthermore, inter-protomer hydrophobic interactions are observed in TM2 and TM10, as depicted in Fig.4, which assists the kink of TM10 upon dimerization. The broad structural changes in these TM helices likely cause the orientation change of Glu563 in the dimer. The increased population of the monomer induced by mutations of Glu563 further supports the view that the region around Glu563 and the dimerization site are structurally linked. It is noteworthy that similar mutation-induced monomer formation was also observed in our earlier work on *Ax*qNOR^10^. The substitution of conserved Glu494 (Glu498 in *Nm*qNOR) to Ala eliminated the hydrogen bond with conserved Asn600 (Asn604 in *Nm*qNOR), leading to the monomer formation. Since this Asn residue is located on TM11 contributing to the dimer formation through the interaction with TM2 in the other protomer, the mutation of Glu494 (Glu498 in *Nm*qNOR) could cause the positional shift of TM11, which affects the dimerization in *Ax*qNOR. Indeed, the low resolution (∼ 5 Å) cryoEM structure of the Glu494Ala mutant of *Ax*qNOR, a monomer, exhibited structural displacements of TM2 and TM10 compared with the wildtype dimer structures of qNOR, from both *A.xylosoxidans* and *N.meningitidis*. (Fig. 8a,b). The displacement of TM10 is broader toward the cytoplasmic end of the helix, where the dimerization site is located. When compared with our *Nm*qNOR wildtype monomer structure, the mutant *Ax*qNOR monomer TM10 also shows a roughly similar conformation, indicating that mutation induced monomerization also produces an altered TM10 property (Fig. 8c). This further emphasizes the importance of dimerization to enable TM10 to adopt an optimal structure for enzyme activity.

**Fig. 8.**
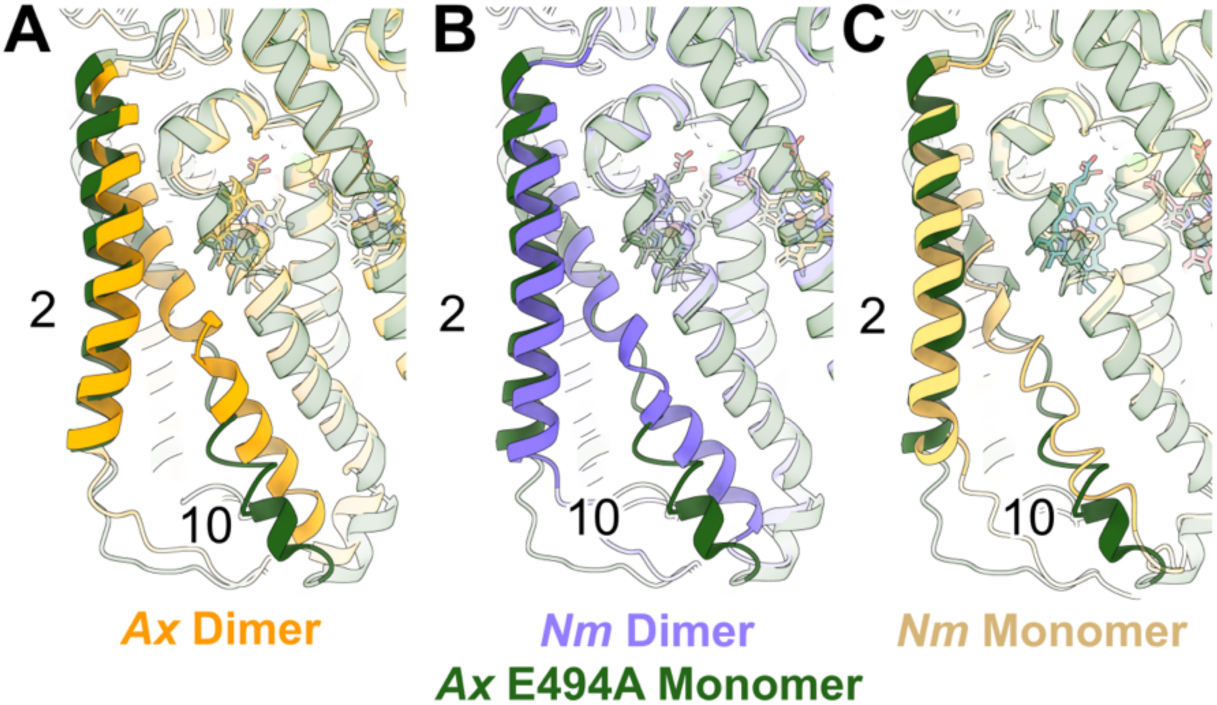
Comparison of transmembrane helical shifts in *Ax*qNOR and *Nm*qNOR monomer-dimer structures. Superposition of the *Ax*qNOR Glu494Ala monomer cryoEM structure (dark green ribbon, PDB ID: 6t6v) with (**A**) the *Ax*qNOR wildtype cryoEM dimer structure (orange ribbon, PDB ID: 8bgw), (**B**) the *Nm*qNOR wildtype cryoEM dimer structure (slate blue ribbon,this study) and, (**C**) the *Nm*qNOR wildtype cryoEM monomer structure (gold ribbon, this study). TM2 and 10 are labeled in all the respective panels and shown as opaquely colored helices, whilst other helices are transparently colored.

The functional rationale of oligomerization in other members of the Heme-Copper Oxidase (HCO) superfamily, such as the oxygen reducing enzyme cytochrome *c* oxidase (C*c*O) is intriguing to compare *Nm*qNOR against. The monomeric C*c*O structure revealed factors that allow for higher oxidase activity versus the dimer^25^. A cholate molecule, which may facilitate dimerization (alongside an array of lipids), was found to disrupt a hydrogen bonding network amongst several water molecules in one proton transfer channel, the K-pathway, which is at the boundary of the dimerization interface. In the C*c*O monomer, this hydrogen-bonded network is intact and is accompanied by a conformational change in Glu62, which is located at the start of the K-pathway (fig. S13). Besides this, no other differences were found between the two forms. This strengthens the idea that a subtle modification in proton uptake at the K-pathway entrance caused the activity differences, which is in stark contrast to qNOR, where large-scale helical rearrangements contributed to side chain conformation changes near the active site. Thus, the mechanism for the enhancement of the catalytic reaction by dimerization in qNOR is unique compared to other HCO enzymes.

In summary, we solved the cryoEM structures of the monomer and the dimer states of *Nm*qNOR with comparably high resolutions. These data indicate that the dimerization-induced broad structural changes at the dimer interface led to the positional shift of the essential Glu563, located near the active site, at the terminal region of the proton transfer channel for an effective NO reduction reaction. The here presented structural and functional information further stresses the importance of the structural determination of each oligomeric state to elucidate the functional properties of the HCO enzymes, including qNOR. Several studies support the critical role of qNOR in survival within host macrophages, producing NO as an immune response, against human pathogens such as *Nm*, *N. gonorrhoeae*^44–46^ and *Staphylococcus aureus*^47^, some of whose multi-drug resistant strains are listed as urgent and serious threats by the World Health Organization^48^. Targeting members of the HCO family in pathogens is seen as a critical junction to help combat antimicrobial resistance^49,50^. Recent work by Nishida and colleagues explored finding a conserved, allosteric inhibitory site in HCO enzymes, (eukaryotic and prokaryotic) including mammalian CcO and bacterial qNOR, to improve the selectivity of inhibitors to reduce unwanted toxicity against host cells. A promising inhibitor, specific to bacterial HCO’s, had a proposed inhibitory mechanism whereby the substrate access channel is partially obstructed^8^. Our current work can also offer an alternate strategy for targeting the dimerization site, such as TM2 and 10, to suppress the enzymatic activity of qNOR upon the dissociation of the functional dimer *in vivo* by chemicals, which would help combat antimicrobial-resistant pathogens.

## Materials & Methods

### Expression and purification of recombinant *Nm*qNOR

The over-expression and purification of *Nm*qNOR fused with BRIL (in this work, *Nm*qNOR represents the one fused with apocytochrome *b*_562_; BRIL) were performed according to previous studies^10^, with minor modifications as follows. The detergent concentration in the buffer (50 mM HEPES pH 8.0, 150 mM NaCl) for the first round of size exclusion chromatography (SEC) using a Superdex 200 26/60 (Cytiva) was adjusted from 0.05 % (w/v) *n*-decyl-β-D-thiomaltoside (DTM; Anatrace) to 0.1 % (w/v) DTM. Fractions corresponding to the dimeric and monomeric forms of *Nm*qNOR from the first round of SEC were separated, pooled, and concentrated to ∼ 10 mg/mL (as judged by the extinction coefficient χ_410_= 213 mM^-1^cm^-1^) using Amicon 50K MWCO centrifugal concentrators (Merck). Dimeric *Nm*qNOR was immediately used to make grids following SEC, whilst monomeric *Nm*qNOR was snap-frozen in liquid nitrogen and stored at −80°C. For the cryoEM analysis of the monomer sample, ∼ 500 µL of 10 mg/mL of sample was thawed and briefly spun down in a centrifuge to remove any large aggregates, before the second round of SEC. The sample was run down a Superdex 200 10/300 Increase (GE Healthcare) column equilibrated in 50 mM HEPES pH 8.0, 150 mM NaCl and 0.1% or 0.05 % (w/v) DTM. Fractions corresponding to the monomer peak were concentrated to ∼ 3 – 15.5 mg/mL (∼ 40 – 170 µM) before grid freezing. Sodium Dodecyl Sulfate Polyacrylamide Gel Electrophoresis (SDS-PAGE) analysis of samples from SEC was performed using NuPAGE 4-12 % Bis-Tris gels, using NuPAGE MES running buffer (Invitrogen). Blue-Native PAGE analysis was also run in parallel according to the manufacturer’s protocol, using NativePAGE 4-16% Bis-Tris gels (Invitrogen). Glu563 variant plasmid DNA was ordered from GenScript (Hong Kong) and transformed into C41 (DE3) cells (Lucigen), before overexpression and purification in a similar fashion to wildtype. The one exception was that SEC was performed in 0.05 % (v/v) *n*-dodecyl-β-D-maltoside (Dojindo), not DTM.

### UV-Visible spectra measurements

UV-Visible spectra of all samples were measured using a U-3900 spectrophotometer (Hitachi). During Nickel-Nitrilotriacetic acid (Ni-NTA) affinity chromatography, fractions were analysed by UV-Visible spectroscopy to determine the A_410_/A_280_ ratio (the so-called Rz value). Fractions with an Rz value ≥ of 0.5 were collected and used for subsequent purification. After size exclusion chromatography, fractions with an Rz value ≥ of 0.7 were concentrated and subsequently used for cryoEM analysis.

### NO reduction activity assay measurement

The NO reduction activity assays were carried out similarly to a previous study^12^. NO reduction rates were calculated from seven independent purifications (five technical replicates) for the dimer and from four independent purifications (five technical replicates) for the monomer samples. NO reduction rates were calculated using the Igor Pro software package (www.wavemetrics.com).

### CryoEM sample preparation and data acquisition

#### *Nm*qNOR_BRIL_ dimer

R1.2/1.3 Cu 300 mesh holey carbon grids (Quantifoil) were glow discharged using a JEC-3000FC Auto Fine Coater (JEOL Ltd.) at 7 Pa, 10 mA for 30 s. Grids were then plunge-frozen in liquid ethane using a Vitrobot Mark IV (Thermo Fisher Scientific) operating at 4°C and 100 % humidity. 3 µL of *Nm*qNOR_BRIL_ (∼ 500 µM) were applied to each grid before blotting for 6 s, using a blot force of 6. Grids were first screened using EPU (Thermo Fisher Scientific) on a Glacios TEM (Thermo Fisher Scientific) and subsequently transferred to a CRYO ARM^TM^ 300 (JEM-Z300FSC; JEOL Ltd.) operating at 300 kV, equipped with an in-column Omega energy filter (20 eV slit width) and K3 direct electron detector (Gatan). SerialEM^51,52^ was used for automated data collection using 5x5 beam-image shift patterns (coma vs image shift calibration was performed before data acquisition). A total of 7,525 movies were collected using a nominal magnification of 60,000x at a pixel size of 0.752 Å/pixel. Each movie was divided into 50 frames using a total dose of 51.2 e^-^/Å^2^ (dose per frame of 1.02 e^-^/Å^2^), with the K3 detector operating in correlated double sampling mode.

#### *Nm*qNOR_BRIL_ monomer

R1.2/1.3 Cu 300 mesh holey carbon grids (Quantifoil) were glow discharged using a JEC-3000FC Auto Fine Coater at 7 Pa, 10 mA for 10 s. Grids were then plunge-frozen in liquid ethane using a Vitrobot Mark IV operating at 8°C and 100 % humidity. 3 µL of *Nm*qNOR-BRIL (initially ranging from 40 – 60 µM) were applied to each grid before blotting for 3 s, using a blot force of 6. Grid-making conditions were optimized over several screening sessions using the Glacios TEM and *EPU* (Thermo Fisher Scientific). Finally, a grid consisting of ∼ 170 µM *Nm*qNOR-BRIL (in 0.05 % DTM, as opposed to 0.1 % DTM), blotted for 3 s using a blot force of 3 exhibited increased particle density with suitable ice thickness for high-resolution data collection. The pre-screened grid was then loaded into a CRYO ARM^TM^ 300 and a total of 8,100 movies were collected using a nominal magnification of 60,000x at a pixel size of 0.752 Å/pixel. Data were collected using 5x5 beam-image shift patterns in SerialEM (coma vs image shift calibration was performed before data acquisition). Each movie (3.66 s exposure time) was divided into 50 frames, using a total dose of 50 e^-^/Å^2^ (dose per frame of 1 e^-^/Å^2^), with the K3 detector operating in correlated double sampling mode. For datasets collected on the CRYO ARM^TM^ 300, fully automated hole centering at medium magnification was performed with the yoneoLocr software, integrated as a SerialEM macro^53^.

### Single particle image processing and 3D reconstruction

#### *Nm*qNOR_BRIL_ dimer

Image processing was performed with RELION-4.0-beta-2-commit-ce2e93^54^, with the workflow shown in fig. S2. 7,525 movies were motion-corrected using RELION’s implementation of MotionCor2 with CTF Estimation performed using CTFFIND-4.1.14. Automated particle picking was performed in crYOLO^55^ and co-ordinate star files were subsequently imported into RELION. 2,574,970 particles were extracted at a box size of 368 pixels and rescaled to 92 pixels (pixel size of 3.008 Å/px). Particles were split into 10 equal subsets of 257,497 particles. After performing 2D classification (VDAM algorithm, K= 150, mask diameter = 185 Å, Limit resolution E^-^ step = 10 Å) on one subset to confirm the initial quality of the images (e.g., no severe orientation bias or degraded particles), 3D classification using a low pass filtered 3D reconstruction from a small dataset (not shown) on a Glacios TEM (Thermo Fisher Scientific) as a reference, was performed on 3 subsets (low pass filter 50 Å, K=4, mask diameter = 190 Å, Limit resolution E^-^ step = 8 Å). ∼365,000 particles were selected and subjected to another round of 3D classification (similar to previous, but with Limit resolution E^-^ step = 5 Å). 221,752 particles were then selected and subject to 3D auto-refinement which reached Nyquist limit (6.14 Å). Particles were extracted to 2.23 Å/px, subject to 3D classification, of which 112,678 particles were 3D-auto-refined to the Nyquist limit (4.5 Å). This cycle of extraction, 3D classification and 3D refinement was continued until 1.09 Å/px pixel size, before CTF Refinement (individual rounds of anisotropic magnification, beamtilt & three-fold astigmatism (trefoil), per-particle defocus & per-micrograph astigmatism), 3D-refinement and postprocessing led to a 2.73 Å (map sharpening B-factor of −66 Å^2^) reconstruction. Subsets 4-6 were then processed similarly to above and the best particles were joined with subsets 1-3 (totaling 326,211 particles). A 3D-refinement and postprocessing job led to a 2.49 Å (−67 Å^2^) reconstruction. CTF refinement (similar fashion as above) and polishing (trained with 10,000 particles and using all frames) were performed before 3D-refinement and postprocessing produced a 2.19 Å (−36 Å^2^) reconstruction.

At this point, a C2 symmetrized map (produced using *relion_align_symmetry* and *relion_image_handler* commands) was used as a reference in a 3D classification job (K=3, mask diameter = 200 Å, Limit resolution E^-^ step = −1). 292,309 particles were carried forward for a further round of 3D classification, with C2 symmetry imposed, leaving 269,127 particles. 3D-refinement and postprocessing lead to a 2.13 Å (−39 Å^2^) reconstruction. CTF refinement (similar fashion as above) and polishing (trained with 10,000 particles and using all movie frames) were performed before 3D-refinement and postprocessing produced a 2.06 Å (−32 Å^2^) reconstruction. Particles were extracted at full pixel size and subjected to two rounds of 3D refinement, CTF refinement (same as above, including four-fold astigmatism (tetrafoil)), and polishing, producing a 1.96 Å (−33 Å^2^) reconstruction (job170). Subsets 7-10 were then processed similarly as described above and the best particles were merged with those from sets 1-6. Two cycles of 3D classification, 3D refinement, CTF refinement, polishing, and 3D refinement led to a final resolution of 1.89 Å (−34 Å^2^) from 357,535 particles.

For the Henderson-Rosenthal plot analysis (fig. S6), particles from the final reconstruction were randomly split into subsets of 500, 1000, 2000, 5000, 10,000, 20,000, 50,000, 75,000,100,000, and 200,000 particles, respectively. Each subset was then refined and sharpened using a soft mask encompassing the whole molecule. The squared value of the attained resolution was plotted against the natural logarithm of each particle subset. The B-factor was calculated by straight-line fitting.

Prior to running 3D Variability Analysis, particle stacks from the final 3D refinement were transferred from RELION to cryoSPARC using UCSF pyEM^56^. The final 3D volume and a custom-made mask to exclude the detergent micelle were imported via the cryoSPARC GUI. 3D Variability Analysis was then performed using the default parameters, aside from a filter resolution of 4 Å. Results were outputted in simple mode using 20 frames and volume down sampling to a 256 pixel box. Motion movies were created in UCSF Chimera v.17.1 using the volume series tool.

#### *Nm*qNOR_BRIL_ monomer

Image processing was performed using cryoSPARC v4.1.1^57,58^, with the workflow shown in fig. S8. All movies were subject to patch-based Motion Correction and patch-based CTF estimation. 104,877 particles were picked from 100 micrographs using the circular blob picker (minimum diameter 50 Å, maximum diameter 150 Å). These were used to generate 2D references for template-based picking against all micrographs, where a total of 4,640,985 particles were picked. 3,963,954 particles were extracted using a 360-pixel box rescaled to 90 pixels (pixel size of 3.008 Å/px). A subset of 150,000 particles were used for ab initio reconstruction (K=3, maximum resolution 7 Å, initial resolution 9 Å). Heterogenous refinement was performed to completion and then ran once again up until one iteration had been completed to generate ‘decoy/bait’ 3D volumes. The remaining particles were subject to two rounds of 2D classification (K=200, maximum alignment resolution 10 Å, maximum resolution 8 Å, force max over poses and shifts = false, mask inner diameter 130 Å, batchsize per class 300, no. of EM iterations = 40 and no. of full iterations = 1) leaving 1,530,505 particles. These were then used for heterogeneous refinement (using the above created ‘good’ and ‘decoy/bait’ classes), and the top two populated classes (795,501 particles) were refined using non-uniform refinement. Particles were re-extracted in a 120 px box (pixel size 2.25 Å/px), before ab initio reconstruction (K= 3, maximum resolution 8 Å, initial resolution 10 Å). Heterogenous refinement was performed with the ab initio models, the previous non-uniform refinement map, and two ‘decoy’ classes from a previous hetero refinement. Two classes (484,196 particles) were subsequently refined and re-extracted in a 180 px box (1.54 Å/px). Ab initio reconstruction and heterogenous refinement with 6 classes led to 2 classes (384,052 particles) which were separately refined (dynamic mask near and far distances of 8 and 20 Å, respectively). The two classes were identical, aside from the handedness which was corrected using the volume tools. Particles were re-extracted to 240 px (1.13 Å/px) and processed similarly as described. For the two best classes (total of 286,686), non-uniform refinement with the integrated fitting of beamtilt, trefoil, anisotropic magnification, and per-particle defocus led to a 2.30 Å reconstruction. After re-extracting in a 324 px box, hetero refinement and non-uniform refinement (similar to above, with fitting of spherical aberration and four-fold astigmatism (tetrafoil)) led to a 2.25 Å (map sharpening B-factor of −64 Å^2^) reconstruction from 274,346 particles. All resolutions were calculated according to the gold standard FSC= 0.143. 3D Variability Analysis was performed using the default parameters, aside from a filter resolution of 4 Å, using a mask excluding the detergent micelle. Results were outputted in simple mode using 20 frames and volume down sampling to a 256 pixel box. Motion movies were created in UCSF Chimera v.17.1 using the volume series tool.

### Model building, refinement and validation

#### *Nm*qNOR_BRIL_ dimer

The starting model (PDB 6l3h) was initially rigidly docked into the final cryoEM map using the UCSF Chimera ‘Fit in Map’ function. The structure was refined using Phenix v.1.15 real_space_refinement module, with the inclusion of restraint files for the heme molecules. Models were manually rebuilt within Coot and subject to further rounds of real space refinement, before validation in MolProbity. Statistics of image processing and refinement are summarized in Supplementary Table 1.

#### *Nm*qNOR_BRIL_ monomer

The starting model (Chain A from the NmqNOR_BRIL_ dimer) was initially rigidly docked into the final cryoEM map using UCSF Chimera ‘Fit in Map’ function and the structure was refined using Phenix v.1.20 real_space_refinement module^59^. To aid initial model building and main chain placement in the disordered regions (residues 245-265 and 567 to 590), the unfiltered cryoEM density map from non-uniform refinement was used. Finer amino acid side chain placement and rotamer fitting were performed using the sharpened map where necessary. Refinements were done using the unfiltered map, with the inclusion of restraint files for the heme molecules. Models were manually rebuilt within Coot^60^ and subject to further rounds of real space refinement, before validation in MolProbity^61^. Statistics of image processing and refinement are summarized in Table S1.

### Figure and graphical illustrations

CryoEM density map depictions and protein model figures were prepared using UCSF ChimeraX v1.4 and v1.6^62^. UCSF Chimera v1.17.3 was used to create the 3DVA movies^63^. Consurf-DB server^64^ and ESpript 3.0^65^ were used to generate sequence conservation and alignment data in fig. S11. Final figure collation was performed within Inkscape 1.1.2 (b8e25be8, 2022-02-05, www.inkscape.org).

## Acknowledgements

We thank Sachiko Hashimoto for culturing of all *Nm*qNOR samples and isolating membrane fractions for purification.

## Author Contributions

HE, MS, CCG, and KF purified and characterised *Nm*qNOR_BRIL_ samples, whilst MS and TT purified and characterized site-directed variants of *Nm*qNOR. CCG and HE froze grids for cryoEM analysis. CG and HS provided overall cryoEM technical support. CCG performed grid screening for dimer and monomer samples, and data collection for the *Nm*qNOR dimer data set. HS performed data collection for the *Nm*qNOR monomer data set. CCG and HE performed image processing. CCG and KM built, refined and validated the atomic models. CCG, YS, KM and TT conceived the study. CCG prepared the figures. CCG, YS, and TT prepared the manuscript.

## Data Availability

Accession codes for the *Nm*qNOR dimer structure are as follows; PDB: 8zgp, EMDB: EMD-60086

Accession codes for the *Nm*qNOR monomer structure are as follows; PDB: 8zgo, EMDB: EMD-60085

## Supplementary Materials for

**Fig. S1.**
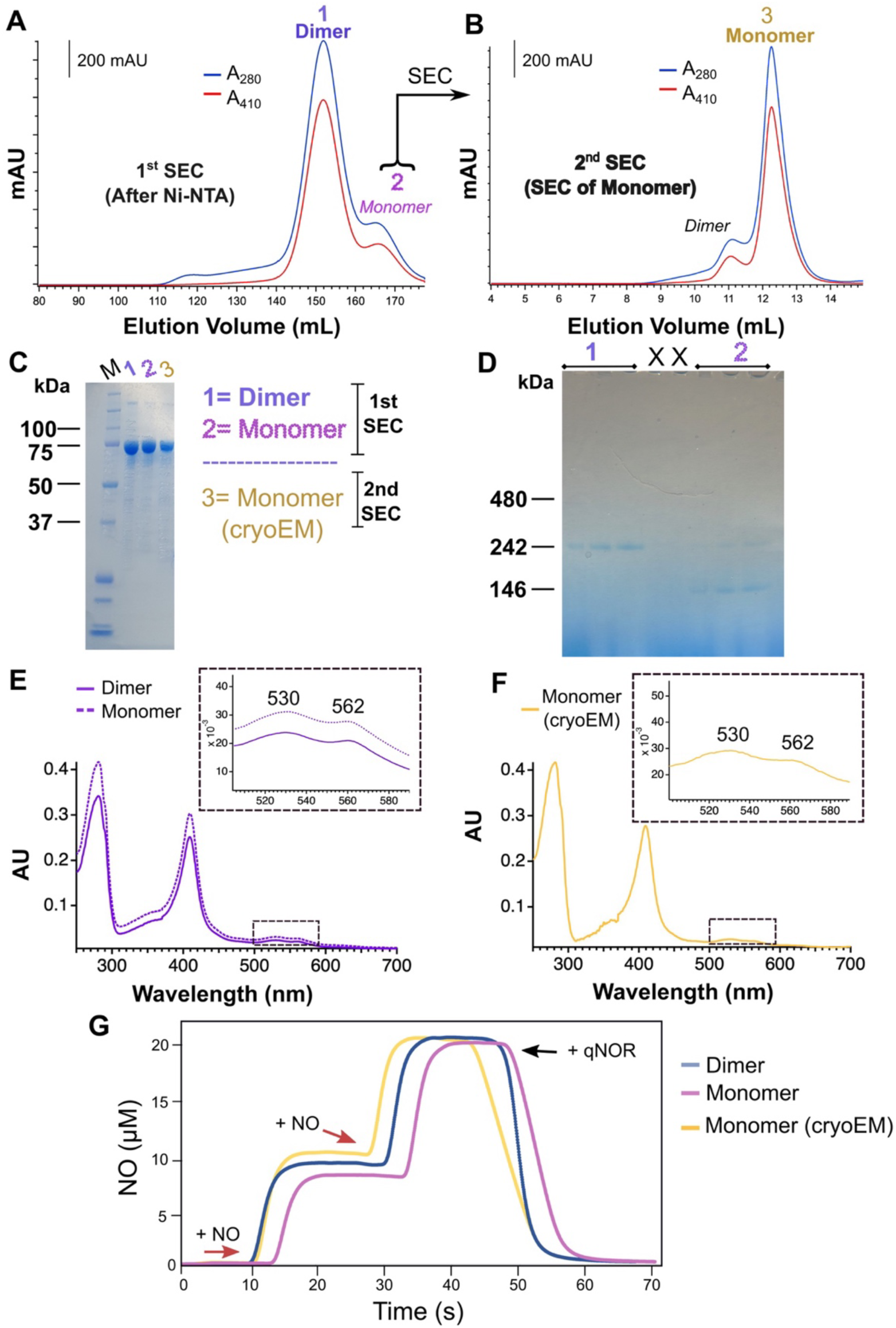
Biochemical characterization of *Nm*qNOR_BRIL_. **A,** Size exclusion chromatogram from the first round of gel filtration i.e., after Ni-NTA purification. Two peaks (peak 1; dimer and peak 2; monomer) are present with peak 1 used for the dimer cryoEM analysis and peak 2 used for a subsequent round of gel filtration shown in (B). **B,** Size exclusion chromatogram of the 1^st^ SEC monomer fractions (termed 2^nd^ SEC). The dominant peak (Monomer) was used for subsequent cryoEM analysis of the monomer qNOR. ‘X’ indicates an empty gel lane. **C,** SDS-PAGE of purified qNOR_BRIL_ samples from 1^st^ SEC and 2^nd^ SEC. Protein markers are indicated on the far left (kDa). **D,** Blue Native PAGE of 1^st^ SEC samples, with labels 1 and 2 corresponding to 1^st^ SEC dimer and monomer peaks, respectively. Protein markers are indicated on the far left (kDa). **E,** UV-Visible spectra of the 1^st^ SEC samples, with peak 1 (dimer) and peak 2 (monomer) drawn as straight and dashed lines, respectively. The boxed region (shown inset) depicts the Q-band region, with peaks at 530 and 562 nm. **F,** UV-Visible spectra of the monomer for cryoEM analysis (2^nd^ SEC sample). The boxed region (shown inset) depicts the Q-band region, with peaks at 530 and 562 nm. **G,** NO consumption assays of wildtype *Nm*qNOR_BRIL_ samples; dimer (blue trace) and monomer (purple trace) from 1^st^ SEC and monomer from 2^nd^ SEC (gold trace). Red and black arrows indicate the addition of substrate (NO) and purified enzyme (qNOR), respectively.

**Fig. S2.**
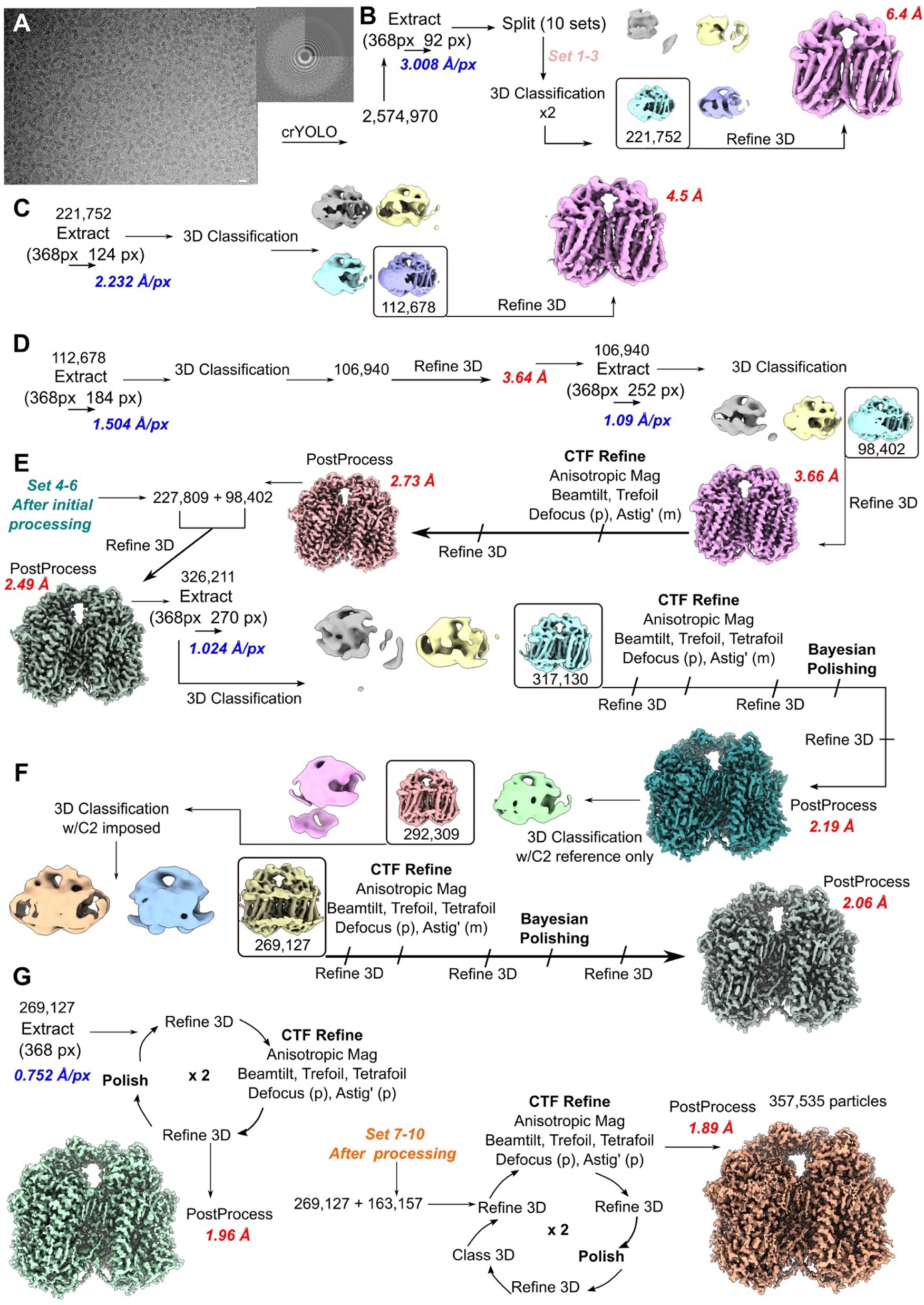
Image processing workflow of the *Nm*qNOR_BRIL_ dimer. **A,** Representative micrograph (left) and associated CTF power spectrum (right)of *Nm*qNOR_BRIL_ dimer. Scale bar = 200 Å. **B,** Particle extraction with x4 binned particles and subsequent processing of subset 1-3. **C,** Particle re-extraction to 2.22 Å/px and subsequent processing of particles. **D,** Particle re-extraction with x2 binned particles and subsequent processing with CTF refinement to 2.73 Å using ∼ 98,000 particles. **E,** Subset 4-6 processed particles and subset 1-3 particles are merged, refined and processed with CTF refinement and polishing till 2.19 Å. **F,** 3D classification with C2 symmetry is performed leaving 269,127 particles to be refined to 2.06 Å.**G,** Particles are re-extracted to unbinned box size and subjected to 2 rounds of 3D refinement-CTF refinement-Polishing, generating a reconstruction below 2 Å. The final set of particles after rough processing (set 7-10) is joined with the present set of particles and subjected to two rounds of 3D refinement-CTF refinement-3D Refinement-Polishing-3D Refinement and 3D classification. The final reconstruction was resolved to 1.89 Å using 357,535 particles.

**Fig. S3.**
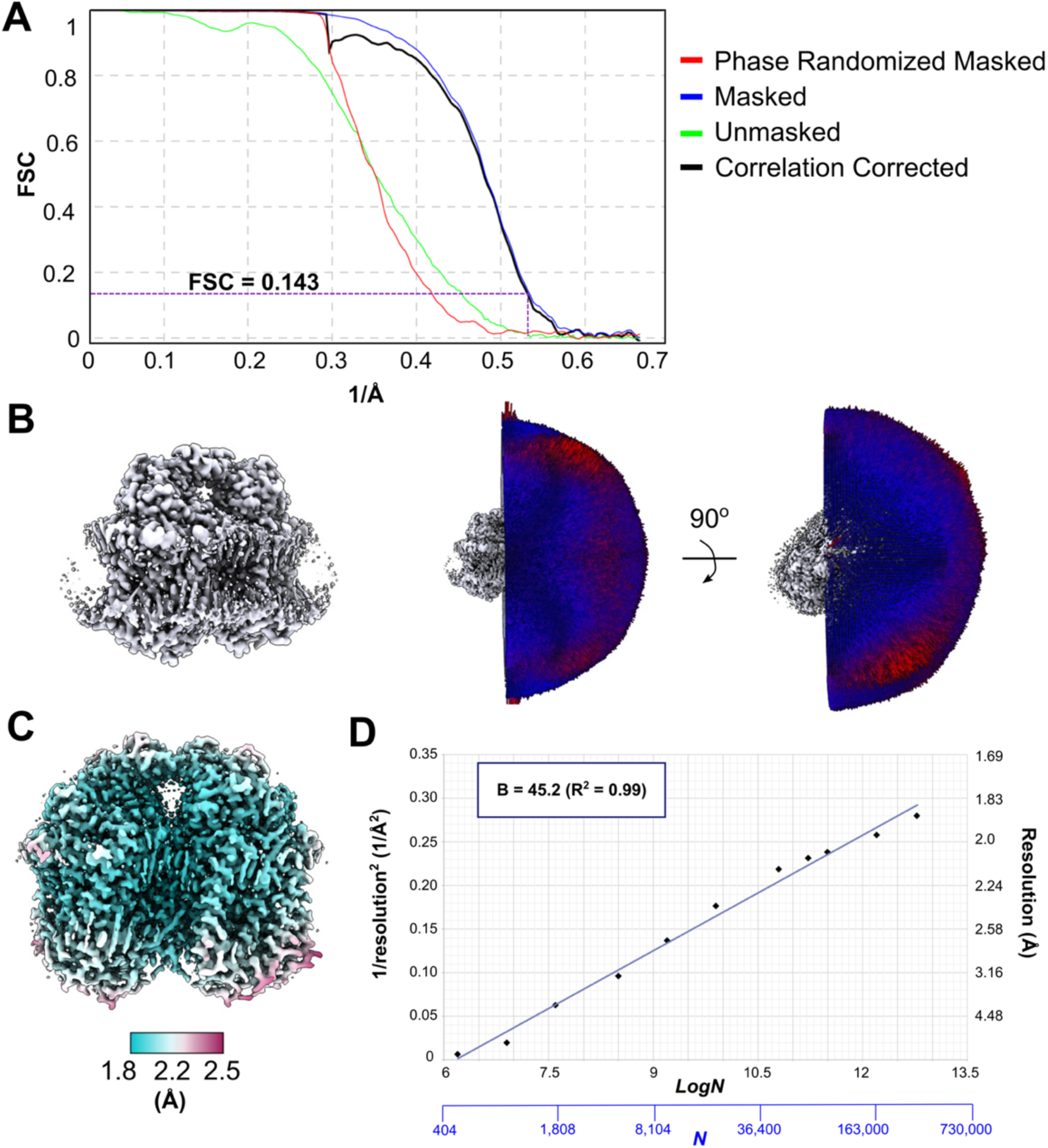
Validation of *Nm*qNOR_BRIL_ dimer cryoEM structure. **A,** RELION generated FSC curves of the *Nm*qNOR_BRIL_ dimer reconstruction. **B,** Euler angle distribution of the C2 symmetrized map. **C,** Local resolution coloring of the unsharpened map, with the resolution color key at the bottom. **D,** Henderson-Rosenthal plot (B-factor analysis) of the *Nm*qNOR_BRIL_ dimer reconstruction. The calculated B-factor from straight line fitting is 45 Å^2^.

**Fig. S4.**
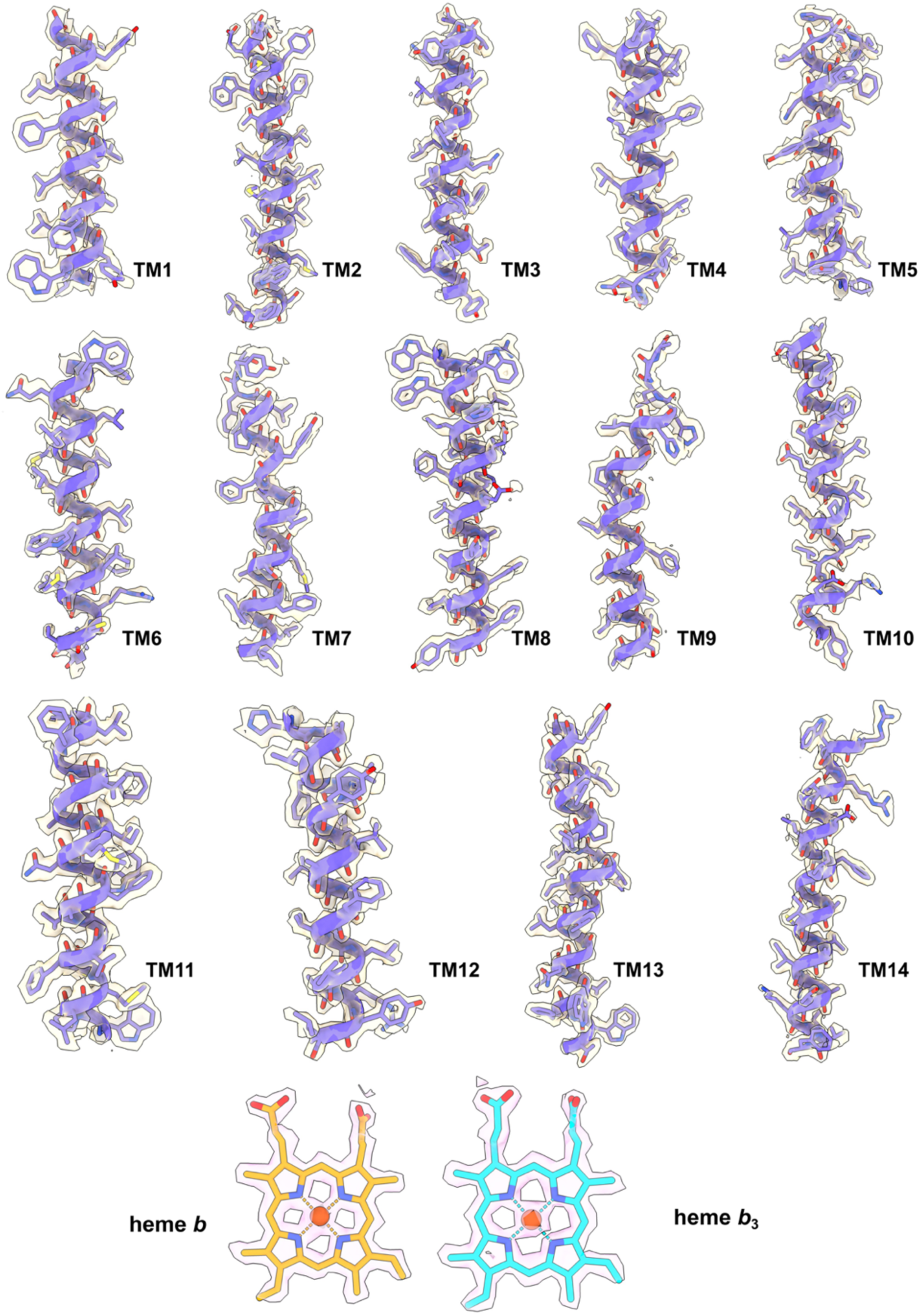
Atomic model and cryoEM density of the *Nm*qNOR_BRIL_ dimer transmembrane helices and heme cofactors. Transmembrane helix (TM) and heme cofactor cryoEM densities with associated models. Sharpened densities (beige) are shown for each TM and heme cofactor.

**Fig. S5.**
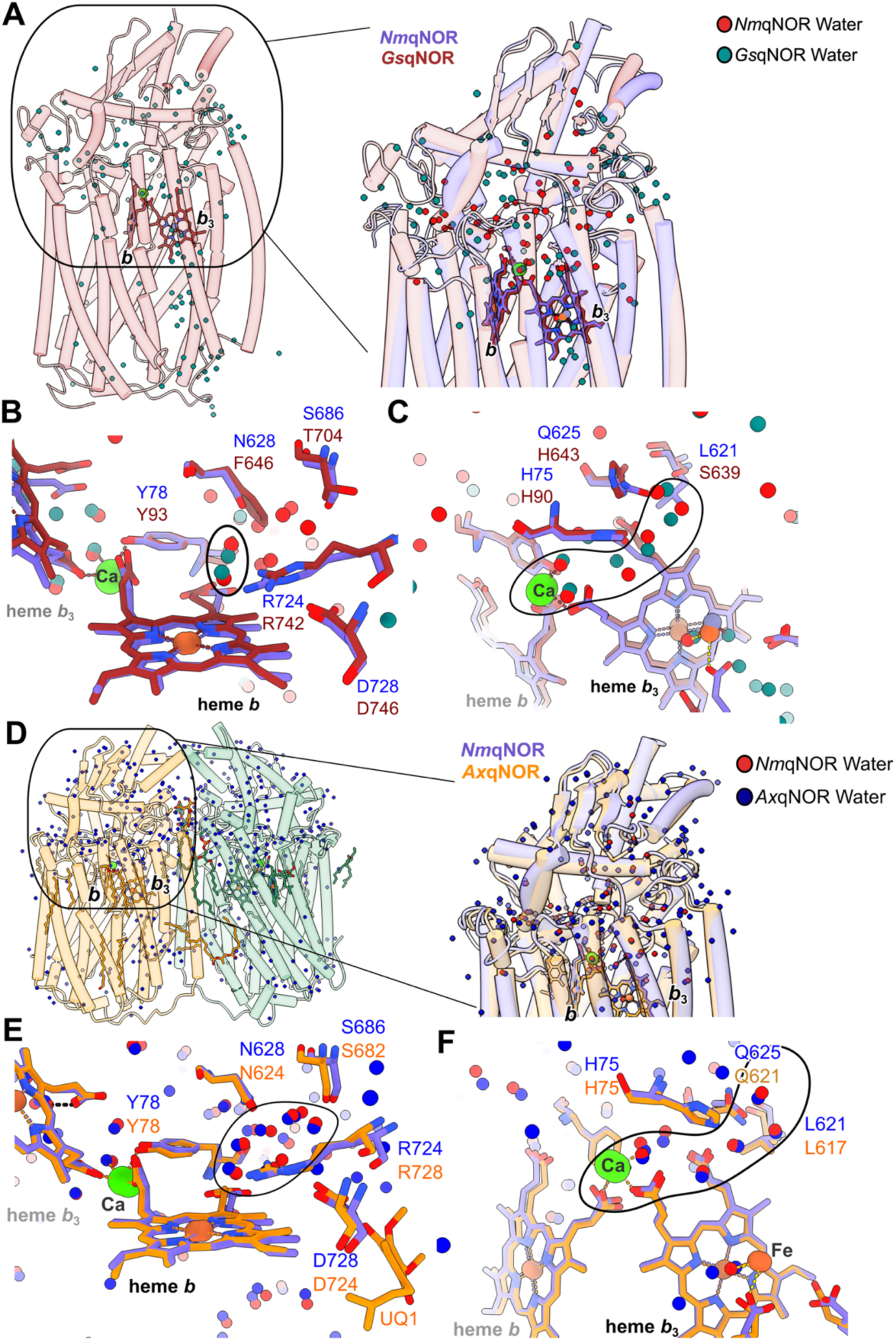
Comparison of the periplasmic water network of *Gs*qNOR, *Ax*qNOR dimer, and the *Nm*qNOR dimer. **A**, Left, structure of *Gs*qNOR in cylinder representation. Right, inset of circled region with one protomer of the *Nm*qNOR dimer structure superposed. Waters for *Gs*qNOR and *Nm*qNOR are coloured as teal and red spheres, respectively. **B,** Comparison of water cluster (black circle) near heme *b* (orange and red sticks) of *Gs*qNOR and *Nm*qNOR. **C,** Comparison of water cluster (black circle) near Ca and heme *b*_3_ of *Gs*qNOR and *Nm*qNOR.**D-F,** Identical to panels a-c, yet, with *Ax*qNOR (orange) water networks (blue spheres) being compared against *Nm*qNOR. PDB ID’s for *Gs*qNOR and *Ax*qNOR are 3ayf and 8bgw, respectively.

**Fig. S6.**
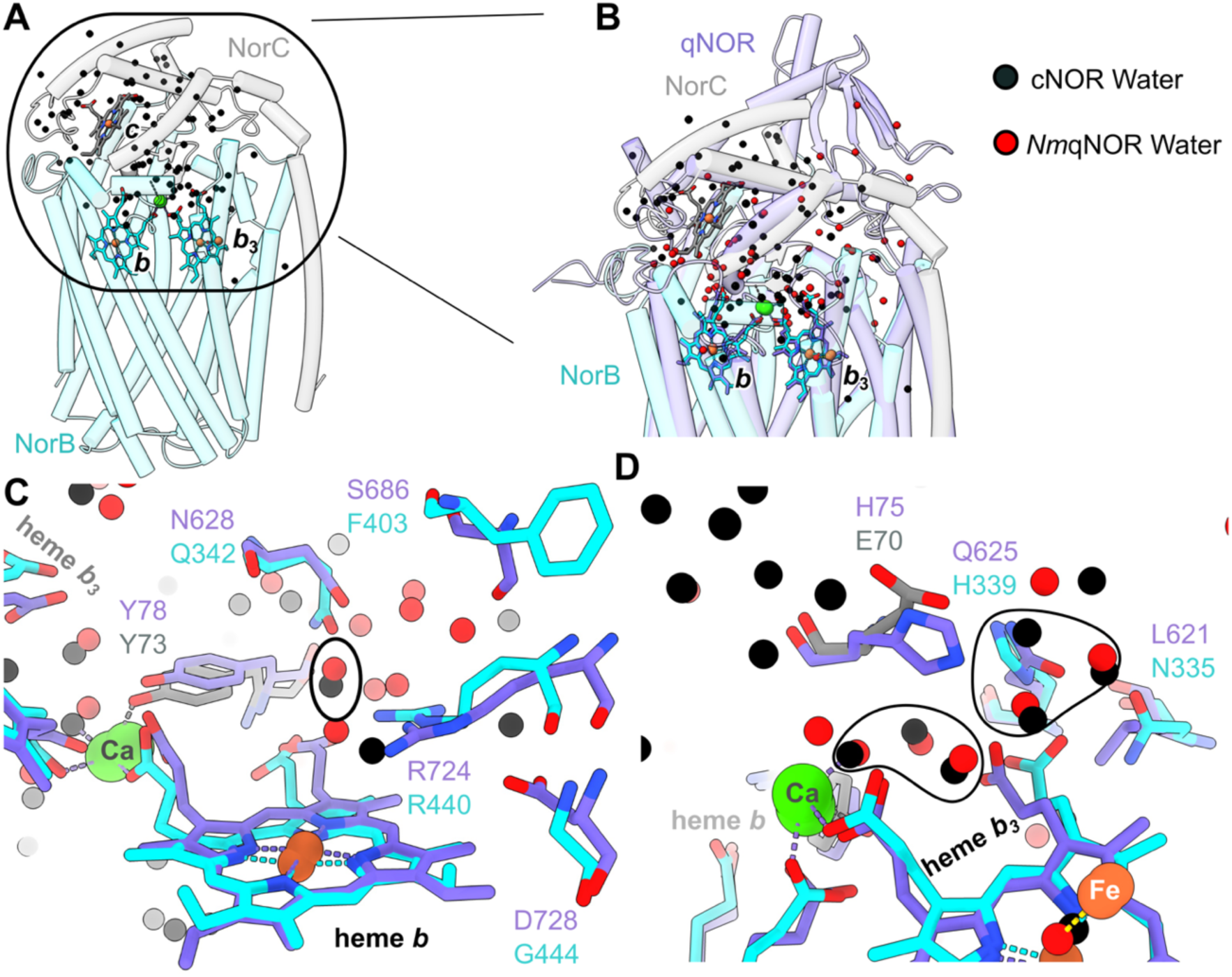
Comparison of the periplasmic water network of *Nm*qNOR dimer and *Pa*cNOR. **A**, Left, structure of *Pa*cNOR in cylinder representation, coloured by subunit (NorC in grey, NorB in cyan). **B,** Inset of circled region with one protomer of the *Nm*qNOR dimer structure superposed. Waters for *Pa*cNOR and *Nm*qNOR are coloured as black and red spheres, respectively. **C,** Comparison of conserved water (black circle) molecules near heme *b*. **D,** Comparison of water cluster (black circle) near Ca and heme *b*_3_ of *Pa*cNOR and *Nm*qNOR.

**Fig. S7.**
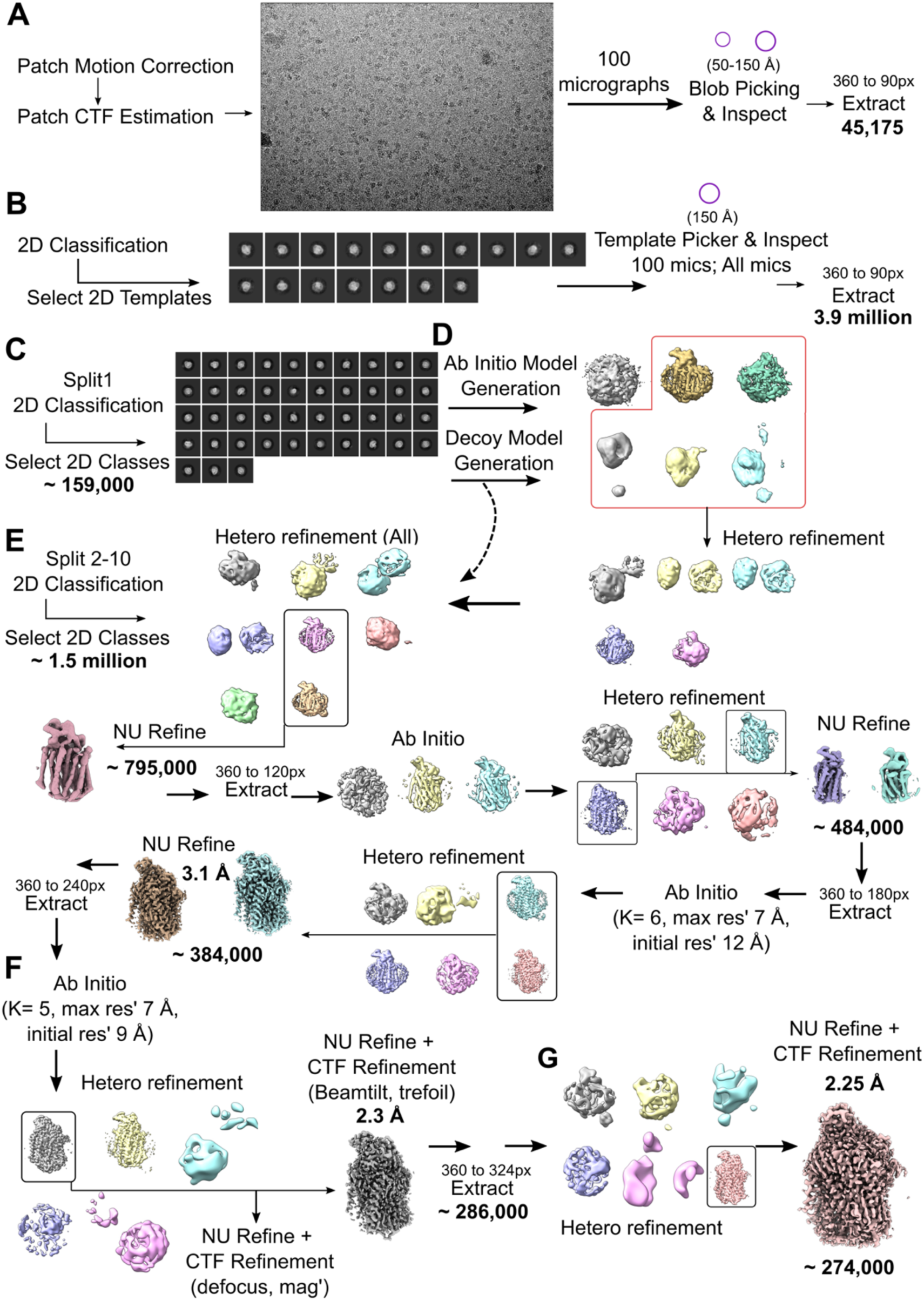
Image processing workflow for the *Nm*qNOR_BRIL_ monomer. **A**, Initial pre-processing of the *Nm*qNOR_BRIL_ monomer dataset. After patch motion correction and CTF estimation, 100 micrographs were randomly selected and initial particle picks were generated using the circular blob picker. Particles were extracted with x4 binning. **B**, 2D template generation for auto picking. After 2D classification, templates were chosen and used for template-based autopicking using 100 micrographs, before applying the optimal parameters to all the micrographs. 3.9 million particles were extracted with x4 binning. **C,**Particle selection for ab initio model generation. 10 % of the picked particles were subjected to 2D classification, with the best class averages used for ab initio model generation. **D**, Particles from (C) were used for ab initio model generation and decoy model (both 3 classes each). **E**, The remaining 90% of particles were subject to 2D classification and the best averages joined with the initial 10 % of the data. Particles were input into a hetero-refinement job using selected step d) classes and the original 3 decoy classes (total of 8 classes). Several rounds of non-uniform refinement (NU Refine), ab initio, hetero-refinement, with intermittent re-scaling of box sizes, led to two 3.1 Å reconstructions. **F**, Ab initio and hetero-refinement were followed by NU Refine with higher order CTF aberration correction to reach 2.3 Å. **G**, A final round of hetero-refinement with a box size of 324 pixels and NU Ref with CTF refinement led to a reconstruction at 2.25 Å.

**Fig. S8.**
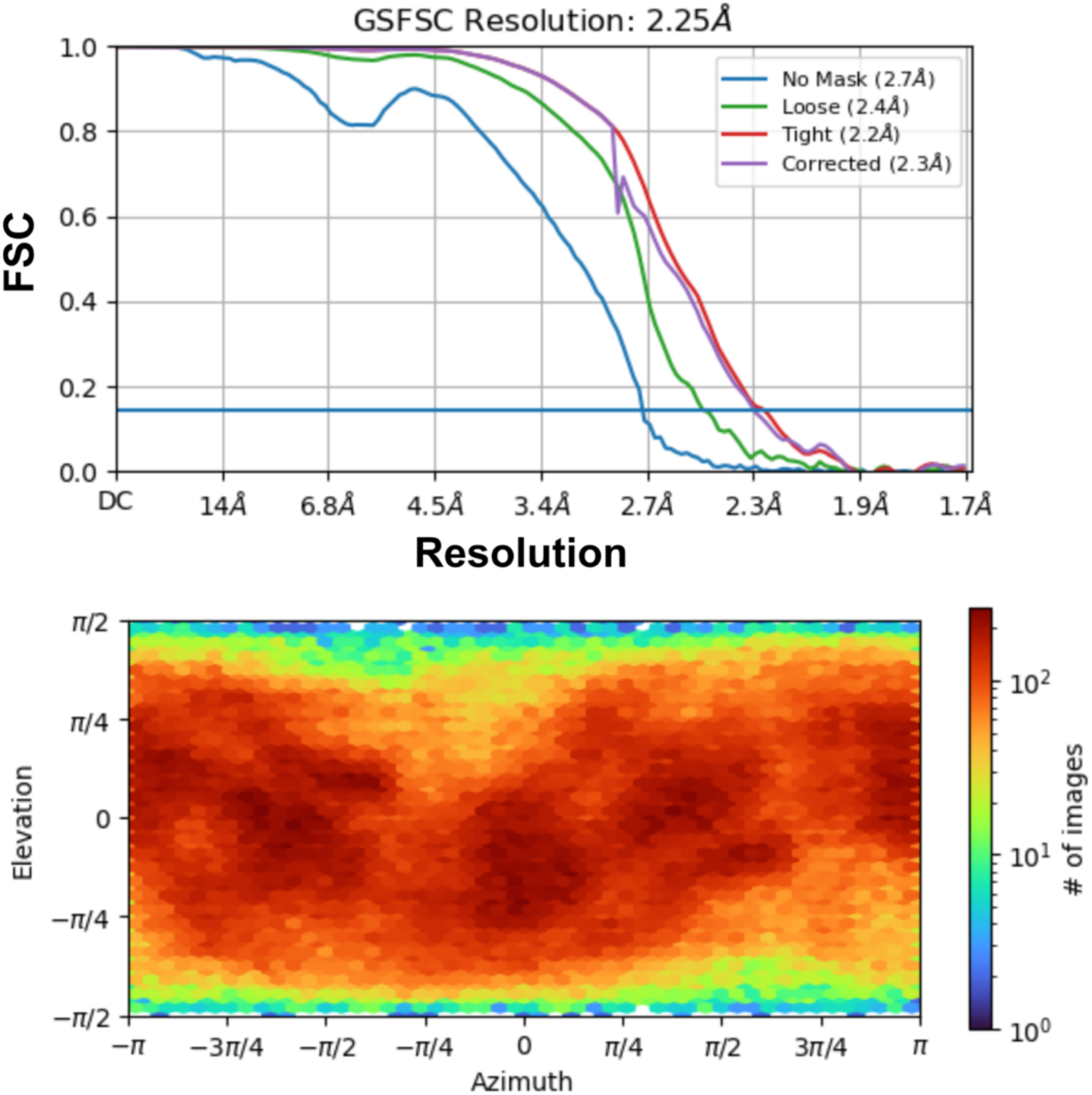
FSC curves and particle orientation plot for the *Nm*qNOR_BRIL_ monomer reconstruction. Top, cryoSPARC generated FSC curves with the horizontal blue line corresponding to FSC = 0.143 (gold standard FSC). Bottom, particle angular distribution plot (bottom) for the *Nm*qNOR_BRIL_ monomer reconstruction,

**Fig. S9.**
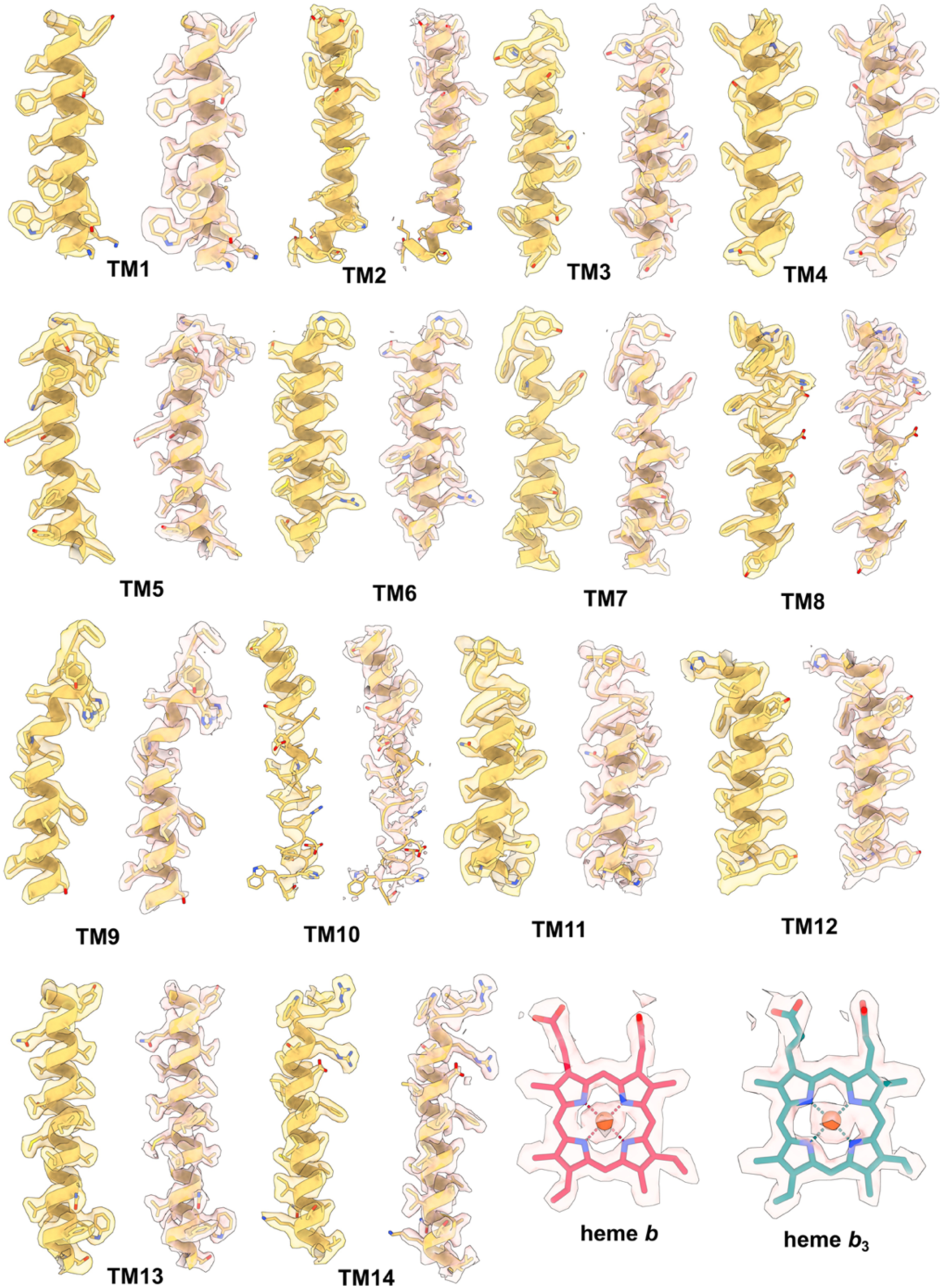
Atomic model and cryoEM density of the *Nm*qNOR_BRIL_ monomer transmembrane helices and heme cofactors. Transmembrane helix (TM) and heme cofactor cryoEM densities with associated models. Both unfiltered map densities (yellow density map, left) and sharpened densities (beige map, right) are shown for each TM. Heme *b* and *b*_3_ densities are from the sharpened map.

**Fig. S10.**
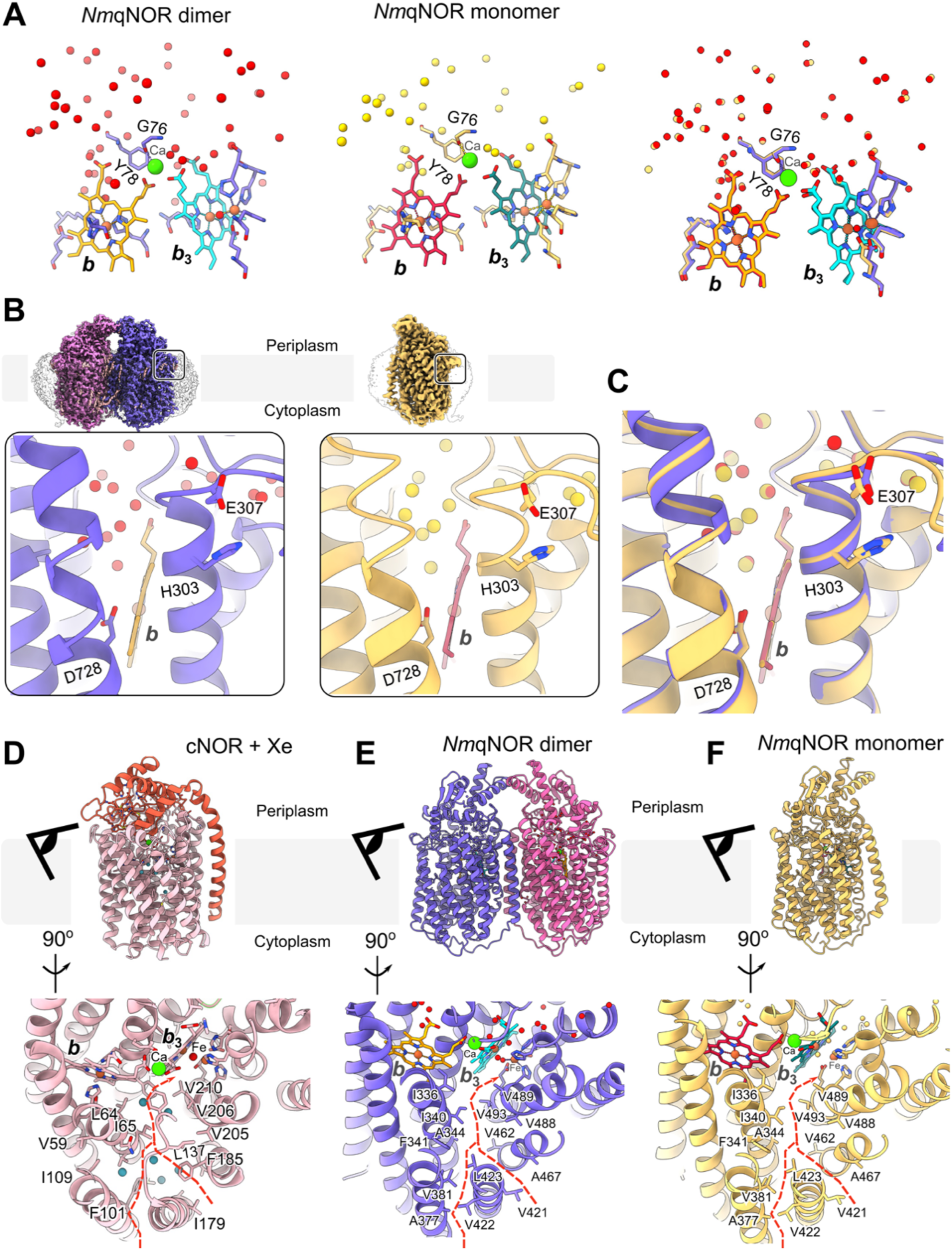
Comparison of *Nm*qNOR dimer and monomer structural elements involved in the enzymatic reaction. **A**, Water molecule distribution around the heme and Ca (green sphere) sites in *Nm*qNOR dimer (left) and monomer (middle). A superposition of each image is shown on the right. **B**, Comparison of the putative quinol binding site in the dimer (left column) and monomer (right column). The inset shows the binding site in more detail, with model (dimer = blue ribbon, monomer = gold ribbon) representation. Water molecules are depicted as red and gold spheres, for the dimer and monomer structures, respectively. **C**, Superposition of the dimer (blue) and monomer (gold) quinol binding sites, with key residues shown at sticks and labeled accordingly. **D**, Potential NO binding and transport channel in *P.aeruginosa* cNOR X-ray crystal structure (PDB ID : 5gux) with xenon (teal spheres). Top, model of cNOR coloured by subunit (NorC in orange and NorB in pink). Bottom, view of the NO binding and transport channel, with residues lining the channel shown as sticks. The red, dashed line indicates the inverted, Y-shaped channel originating from within the membrane that could transport NO toward the catalytic heme *b*_3_ – Fe site. **E-F**, The same as (D), except for the *Nm*qNOR dimer and monomer cryoEM structures, respectively.

**Fig. S11.**
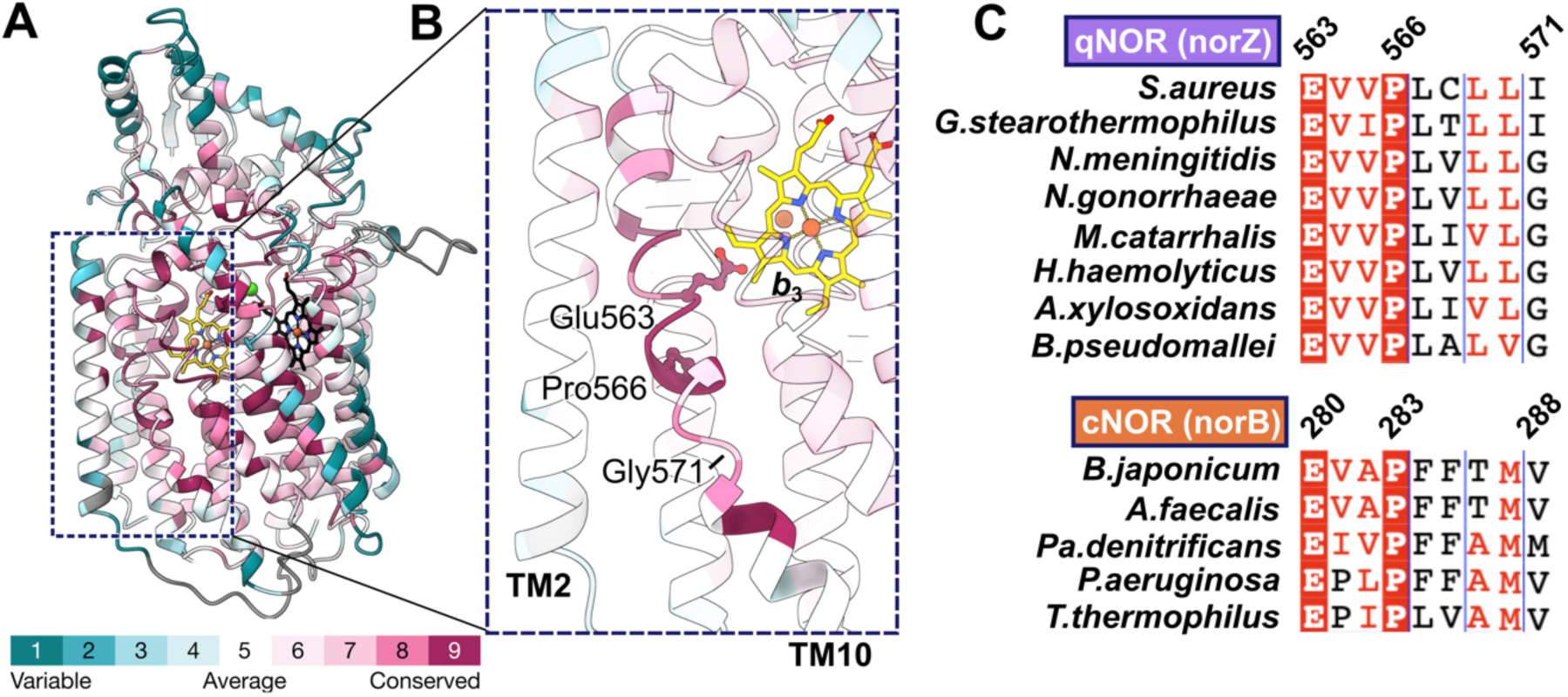
Amino acid conservation profile of the TM10 kink-turn region in bacterial NORs. **A,** *Nm*qNOR (chain A of the dimer) colored by ConSurf-DB conservation; teal (grade 1) indicates residues with more variable sequence through till maroon (grade 9), indicating highly conserved sequence position. **B,** Zoomed in view of the dashed box from (A), depicting the TM10 kink-turn region generated by Pro566 and Gly571, and Glu563 relative position. **C,** Sequence alignment of the NOR (top, qNOR norZ gene and bottom, cNOR norB gene) TM10 kink-turn region from selected organisms, with residues at position 563, 566 and 571 labeled. Figure was generated using the ClustalW and ESpript 3.0 servers.

**Fig. S12.**
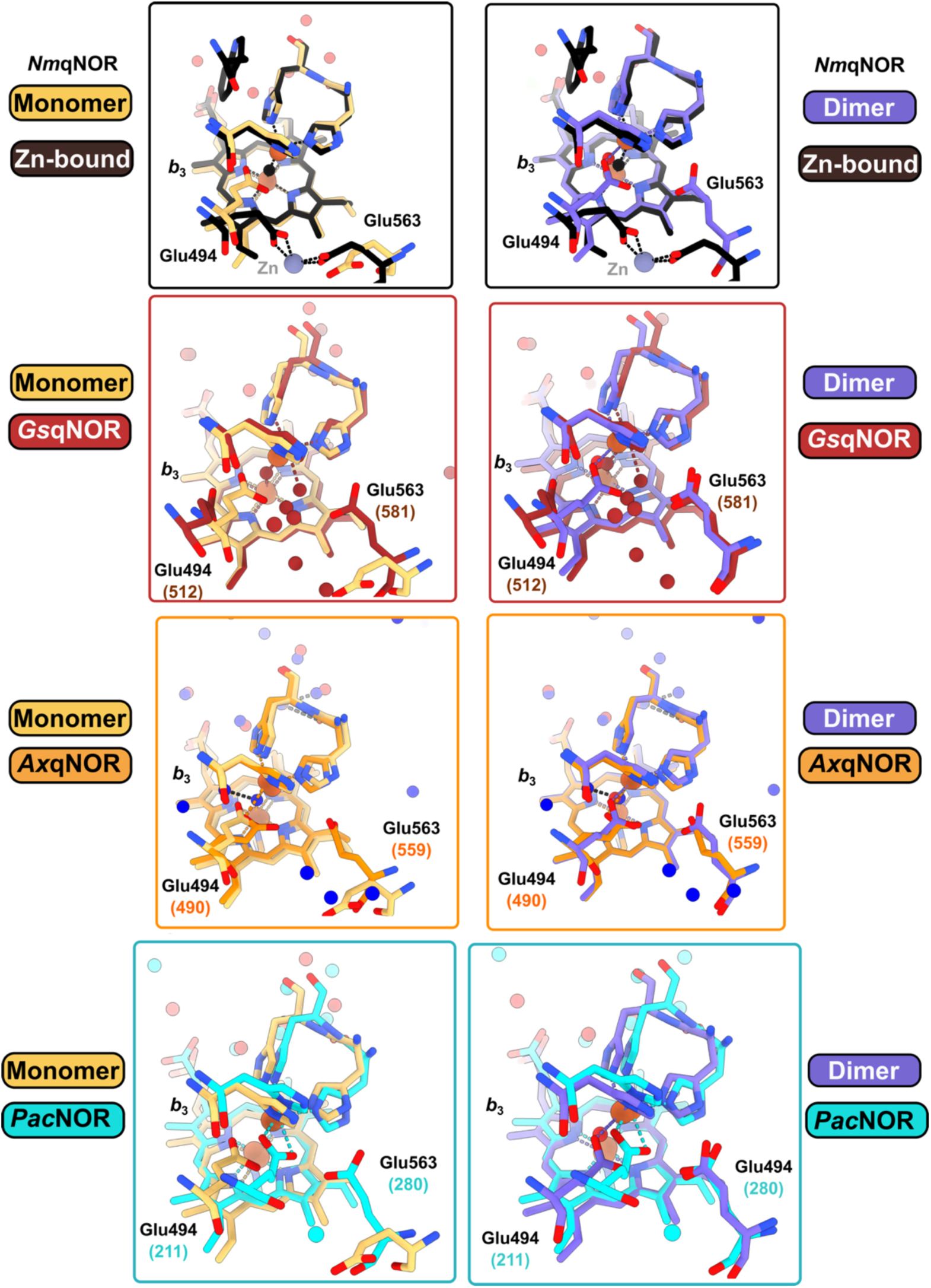
Comparison of active site glutamate conformations amongst NOR’s. Superposition of *Nm*qNOR dimer (slate blue sticks, right hand column) and monomer (gold sticks left hand column) active site against the *Nm*qNOR Zn-bound X-ray crystal structure (black sticks, PDB ID: 6l1x), *Gs*qNOR X-ray crystal structure (brown sticks, PDB ID: 3ayf), *Ax*qNOR dimer cryoEM structure (orange sticks, PDB ID:8bgw) and *Pa*cNOR X-ray crystal structure (cyan sticks, PDB ID: 3o0r) respective active sites. Residue labels in parentheses are numberings for respective protein molecules compared to that of *Nm*qNOR.

**Fig. S13.**
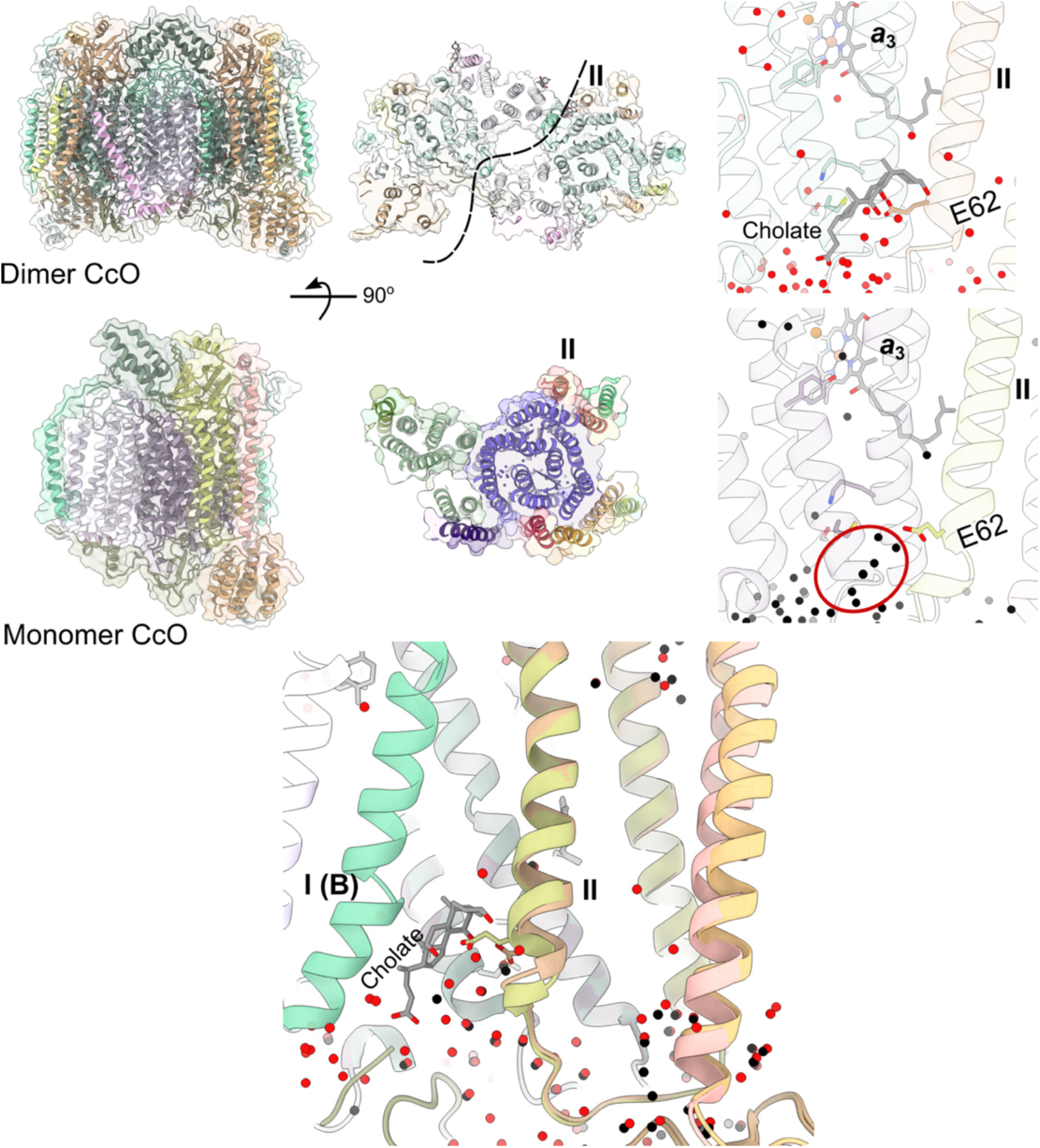
Bovine cytochrome *c* oxidase monomer-dimer comparison. Left, ribbon and surface models for bovine dimer C*c*O (top, PDB: 2dyr) and monomer (middle, PDB : 6jy3) C*c*O, with rotated views shown in the middle. The dimer separation is shown with a dashed black line. Right, view of the entrance of the K-pathway for both dimer (top panel) and monomer C*c*O (bottom panel). The dimer contains a cholate (grey stick) between subunit II of one monomer and subunit I of the other. Monomer C*c*O is free of cholate and contains a chain of hydrogen bonded waters (black spheres, circled in red) with E62 helping co-ordinate them. The bottom panel shows a superposition of both dimer and monomer CcO, with water molecules of dimer and monomer C*c*O shown as red and black spheres, respectively. Subunit I of the dimer (I (B)) and the cholate molecule form the dimerization interface with subunit II.

**Table S1.** CryoEM data collection and image processing.

|  | *Nmq*NOR dimer<br>(EMDB : EMD-60086, PDB : 8zgp) | *Nmq*NOR monomer<br>(EMDB : EMD-60085, PDB : 8zgo) |
| --- | --- | --- |
| **<u>Data collection parameters</u>** |  |  |
| **Microscope** | CRYO ARM™ 300<br>(JEOL Ltd.) |  |
| **Acceleration Voltage (kV)** | 300 |  |
| **Detector\*** | K3 Direct Electron Detector<br>(Gatan) |  |
| **Nominal magnification** | 60,000x |  |
| **C2 aperture diameter (μm)** | 50 |  |
| **In-column Omega energy**<br>**filter slit width (eV)** | 20 |  |
| **Pixel size (Å/pixel)** | 0.752 |  |
| **Target defocus range**<br>**(μm)** | -1.2 to -1.6 |  |
| **Data Acquisition Software** | SerialEM |  |
| **Exposure time (s)** | 2.26 | 3.66 |
| **Total exposure (e<sup>-</sup>/Å<sup>2</sup>)** | 51.19 | 50 |
| **Movie frames** | 50 |  |
| **Exposure per frame**<br>**(e<sup>-</sup>/Å<sup>2</sup>/frame)** | 1.02 | 1.00 |
| **<u>Image Processing</u>** |  |  |
| **No. of movies** | 7,525 | 8,100 |
| **Particle extraction box size (pixels)** | 368 | 384 |
| **Final particle box size (pixels)** | “ | 324 |
| **Number of extracted particles (post particle picking)** | 2,574,970 | 3,953,954 |
| **Particles in final 3D reconstruction** | 357,535 | 274,346 |
| **Reconstruction Symmetry** | C2 | C1 |
| **Map sharpening B-factor (Å<sup>2</sup>)** | - 32 | - 64 |
| **Map Resolution (Å) (FSC threshold= 0.143)** | 1.89 | 2.25 |
| **Starting model** | PDB 6I3h | Chain A from *NmqNOR* dimer (this work, PDB 8zgp) |
| **CC (mask)** | 0.84 | 0.87 |
| **CC (main chain)** | 0.83 | 0.84 |
| **CC (side chain)** | 0.80 | 0.81 |
| **CC (ligands)** | 0.83 | 0.76 |
| **Non-hydrogen atoms** | 12,156 | 5,981 |
| **Protein residues** | 1,484 | 741 |
| **Ligands** | 8 | 4 |
| **Mean B-factor (protein) (Å<sup>2</sup>)** | 29.38 | 95.72 |
| **Mean B-factor (ligand) (Å<sup>2</sup>)** | 15.82 | 77.24 |
| **R.M.S.D. bond lengths (Å)** | 0.005 | 0.003 |
| **R.M.S.D. bond angles (°)** | 0.717 | 1.32 |
| **Clash score** | 5.41 | 7 |
| **Rotamer outliers (%)** | 1.0 | 3.8 |
| **MolProbity score** | 1.43 | 2.01 |
| **Ramachandran plot**<br>**favored (%)** | 97.56 | 96.74 |
| **Ramachandran plot**<br>**allowed (%)** | 2.44 | 3.26 |
| **Ramachandran plot**<br>**outliers (%)** | 0 | 0 |

\* operated in correlated double sampling (CDS) mode

**Movie S1.**
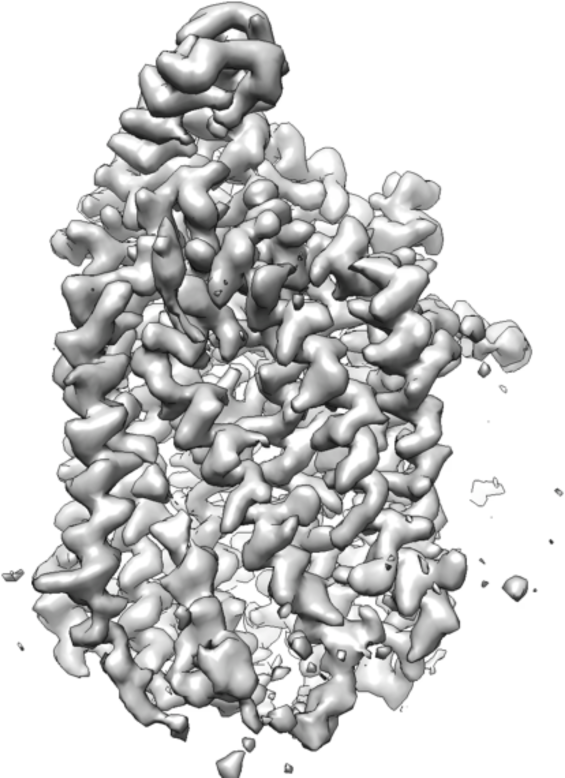
Component 1 of *Nm*qNOR monomer 3DVA.

**Movie S2.**
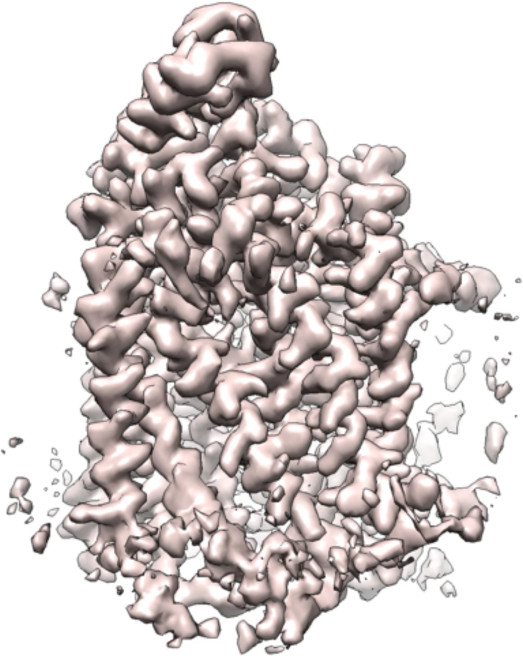
Component 2 of *Nm*qNOR monomer 3DVA.

**Movie S3.**
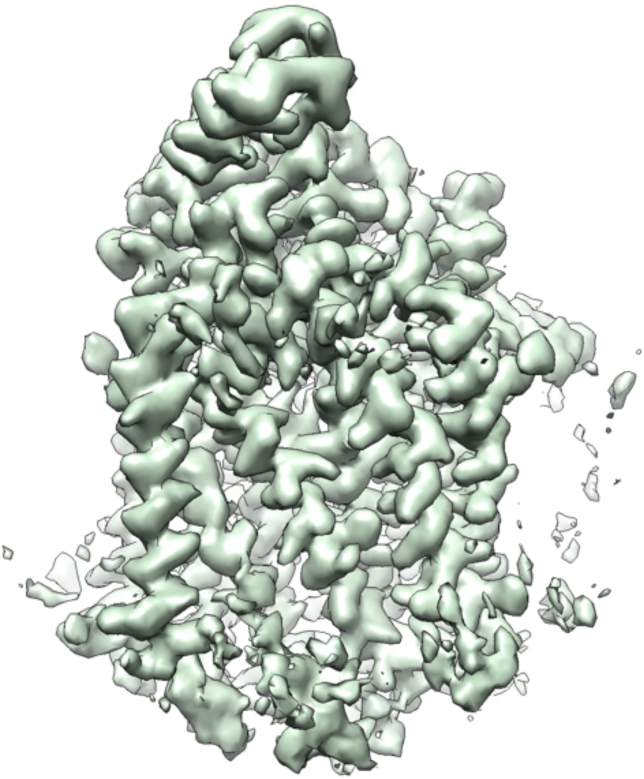
Component 3 of *Nm*qNOR monomer 3DVA.

**Movie S4.**
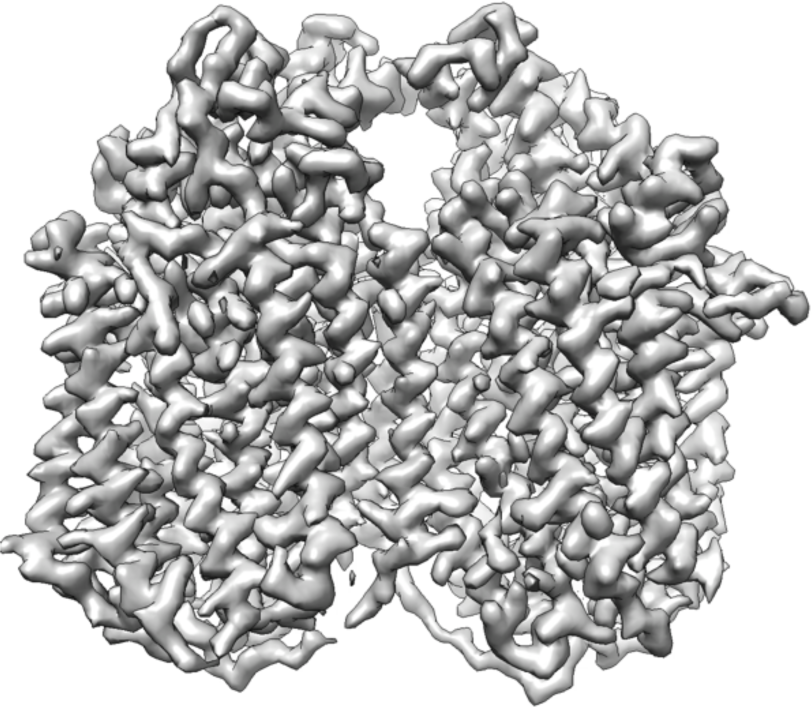
Component 1 of *Nm*qNOR dimer 3DVA.

**Movie S5.**
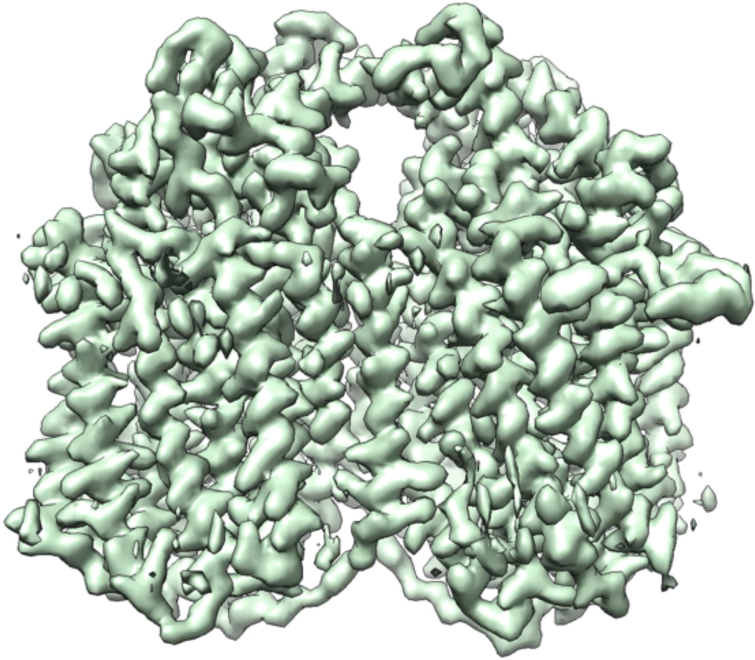
Component 2 of *Nm*qNOR dimer 3DVA.

**Movie S6.**
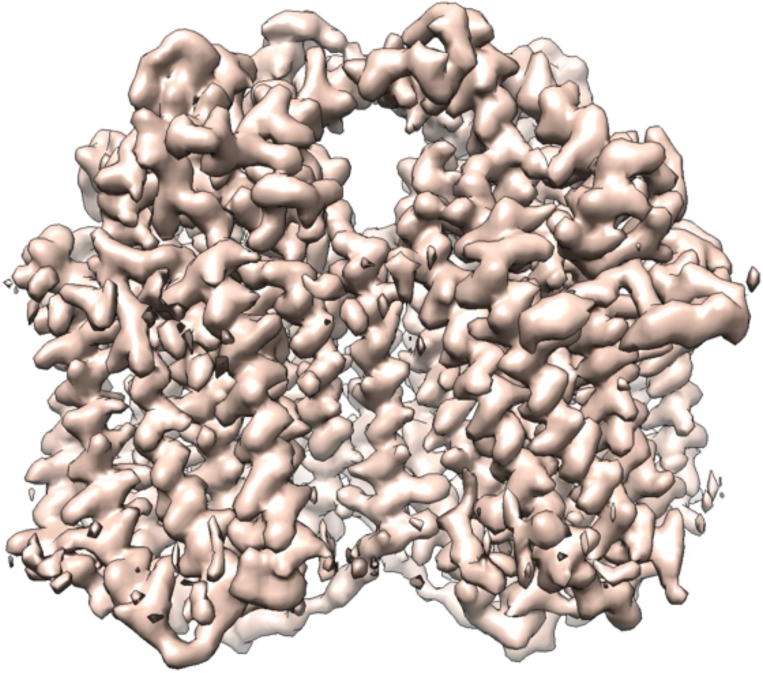
Component 3 of *Nm*qNOR dimer 3DVA.

